# Biomolecular condensates bridge experiment and theory of mass-conserving reaction-diffusion systems in phase separation

**DOI:** 10.1101/2024.08.08.607271

**Authors:** Cheng Li, Man-Ting Guo, Xiaoqing He, Quan-Xing Liu, Zhi Qi

## Abstract

Phase separation is crucial in biological processes and ecological resilience, yet experimental evidence supporting validity of mass-conserving reaction-diffusion (MCRD) models in describing phase separation remains rare. Here, we identified one type of biomolecular condensates – double-stranded DNA (dsDNA) and the human transcription factor p53 as forming dsDNA-protein interactive co-condensates (DPICs) as an experimental model, where dynamic DPICs evolve into droplet-like patterns through reversible autocatalytic biochemical reactions. Thus, we can use the MCRD model to describe this experimental system. The pattern formation depends on concentration of protein and dsDNA and their local nonlinear interactions, which were experimentally tested at meso-scale and integrated into an MCRD model. Our results provide direct evidence that the experimental data-driven MCRD models can reproduce the observed phase diagram and scaling-invariance coarsening patterns, thus, providing a compelling mechanism for phase-separation pattern in other systems.

## Introduction

Pattern formation is a fundamental process in biological development and ecosystem functioning, occurring across scales from the micrometer level, such as cytoskeletal structures (*1, 2*), to centimeter-scale zebrafish stripes (*3, 4*), and extending to kilometer-scale tiger bush vegetation patterns (*5, 6*). These patterns arise from localized, nonlinear interactions among system components and/or their environment. Both model predictions and experimental evidence suggest that self-organized patterns are critical for regulating biological signal processing (*7, 8*) and enhancing ecological resilience to environmental change (*6, 9*). Research in this field predominantly combines observational studies with mathematical modeling (*3, 5, 10, 11*) or experimentally investigates the interactions underpinning scale-dependent feedback mechanisms (*12, 13*).

Among these pattern formations, one important type is phase separation. Recent studies in macroscale pattern formation, spanning biogeomorphology and microbial ecology (*14-16*), have identified phase separation as a key driver in systems such as worm blobs (*17*), tidal flat mussel beds (*18*), and patterned ground (*19*). Experimental and theoretical investigations of phase separation—often characterized by coarsening behaviors (*9, 17, 19, 20*)—have been extensively conducted in physical (*21, 22*), chemical (*23, 24*), biomolecular (*25-29*), and biogeomorphological (*9, 20*) systems. Despite these progresses, studies that directly bridge experimental observations and theoretical models in phase separation remain rare (*20, 27, 30*).

Over the past decades, numerous theoretical models have been developed to elucidate the mechanisms underlying phase separation phenomena. The Cahn–Hilliard model, along with its coupling to kinetic dynamics, has been widely employed to replicate coarsening behaviors observed in non-equilibrium thermodynamic systems (*24, 25, 27, 31-34*). In the context of active particle systems, motility-induced phase separation (MIPS) has emerged as a robust theoretical framework, offering insights into the spontaneous self-organization of active particles into distinct high- and low-density phases (*20, 35, 36*). Beyond a large class of phase separation, an alternative approach using reaction-diffusion models based on reversible autocatalytic chemical reactions has been developed to model coherent localized structures (*9, 37, 38*).

We test the hypothesis that reaction-diffusion models alone can explore the phase-separation patterns in biomolecular condensates. Within living cells, macromolecules such as nucleic acids and proteins are known to form mesoscale biomolecular condensates through liquid-liquid phase separation (LLPS) (*39, 40*). LLPS plays a critical role in various biological functions (*7, 41, 42*). However, its dysregulation is implicated in pathological conditions, including neurodegenerative diseases and cancer (*43*).

We focused on a specific phase-separation phenomenon in dsDNA-protein interactive co-condensates (DPICs), where neither dsDNA nor protein alone can undergo phase separation. Instead, DPIC formation requires the protein to associate with multiple distinct dsDNA molecules, creating a network-like architecture in solution. Examples include systems such as cGAS-dsDNA (*44*), VRN1-dsDNA (*45*), and the KLF4 DNA-binding domain-dsDNA interaction (*46*). Interestingly, we recently discovered that a truncated version of the human transcription factor p53, when combined with dsDNA, can also form DPICs (*47*).

The DPIC formation can be represented as a reversible autocatalytic biochemical reaction, as depicted in Fig. 1A: DPIC (containing *m* protein molecules and dsDNA) + free protein ⇆ DPIC (containing *m* + 1 protein molecules and dsDNA). For a broad class of phase-separation phenomena, reaction-diffusion models based on reversible autocatalytic chemical reactions have been employed to describe coherent, localized structures (*37*). Given that the free energy change (ΔG) of the protein-dsDNA binding reaction is typically negative (ΔG < 0) (*48*), the autocatalytic reaction is thermodynamically favorable and will progress spontaneously toward equilibrium, maintaining detailed balance. Additionally, under *in vitro* experimental conditions, the total mass of substrates remains conserved during DPIC formation. Therefore, we employed the mass-conserving reaction-diffusion (MCRD) model to investigate the potential mechanisms underlying the formation of these biomolecular condensates.

**Fig. 1.**
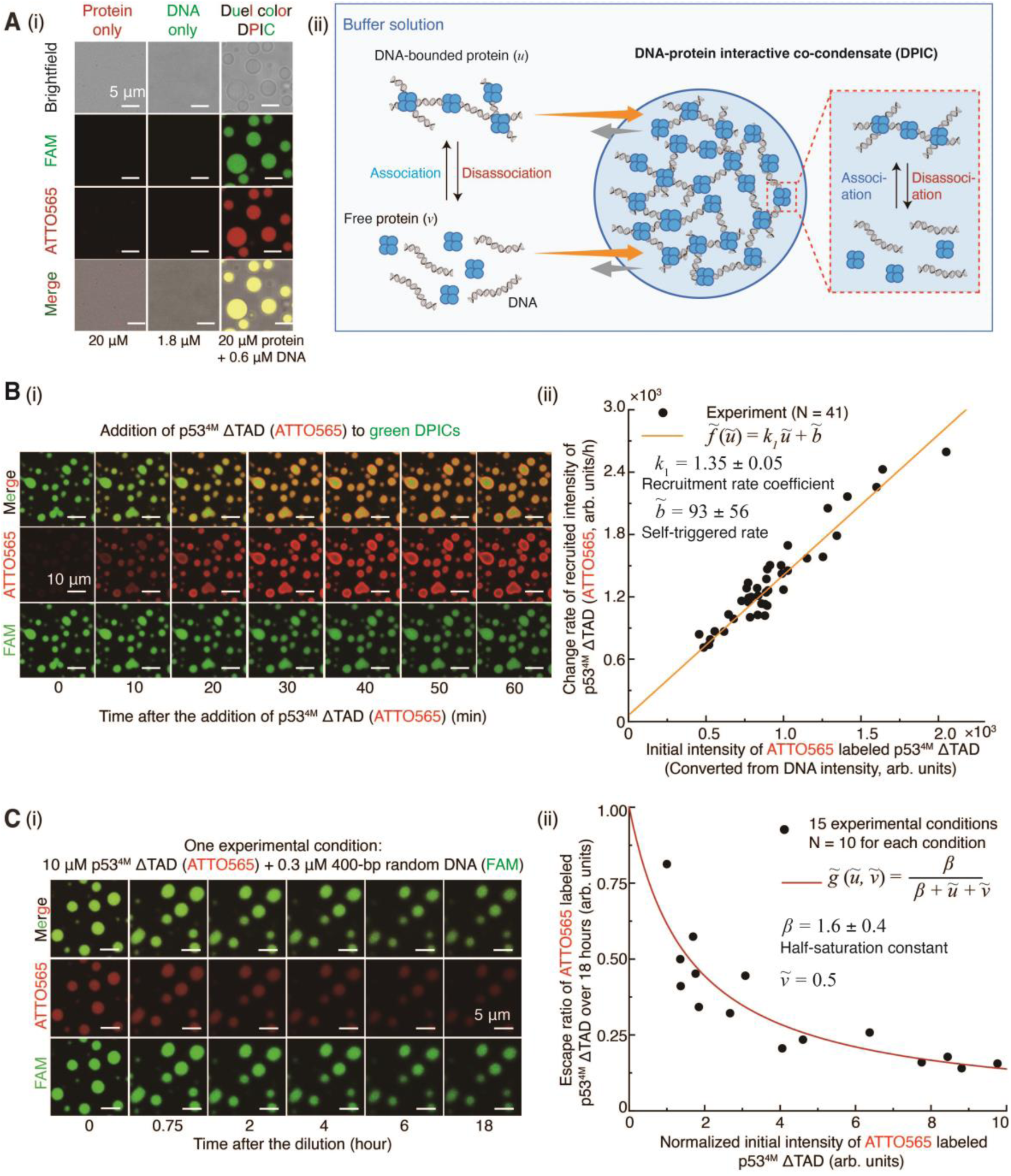
DPICs exhibit both positive and negative local concentration-dependent dynamic behaviors of proteins. (**A**) (i) *In vitro* droplet assays. From left to right: 20 μM p53^4M^ ΔTAD labeled by ATTO565, 1.8 μM 400-bp random dsDNA labeled by FAM, DPICs involving the mixing of 0.6 μM 400-bp random dsDNA labeled by FAM with 20 μM p53^4M^ ΔTAD labeled by ATTO565. (ii) Schematic diagram of the formation process of DPIC. In this picture, *v* represents free protein and *u* represents dsDNA-bound protein. The yellow arrows indicate the recruitment of proteins by DPIC, and the gray arrows indicate the escape of proteins from DPIC. (**B**) (i) Time course of the recruitment of free protein into the two color-labeled DPICs, with time indicated in minutes. (ii) The rate of change in the average intensity of recruited ATTO565-labeled p53^4M^ ΔTAD at equilibrium versus the initial amount of p53^4M^ ΔTAD inside the DPIC (Supplementary Fig. 4B(v) and **Methods**) was plotted, which can be fitted by a recruitment rate, 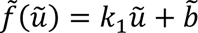. (**C**) (i) Time course of the escaped protein from the two color-labeled DPICs, with time indicated in minutes. Here is one specific experimental condition of 10 µM p53^4M^ ΔTAD labeled with ATTO565 and 0.3 µM 400-bp random dsDNA labeled with FAM. All data of 15 different experimental conditions were shown in Supplementary Fig. 5. (ii) The escape ratio of proteins at the 18-hour time point versus the normalized initial amount of p53^4M^ ΔTAD inside DPIC at 0-hour time point was plotted, which can be fitted by an escape rate, 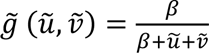. Independent *in vitro* droplet experiments in A, B(i), and C(i) were conducted three times (n = 3). It is pertinent to note that in the subsequent sections, mathematical symbols with a tilde denote the fluorescent intensity measured from experiments, whereas symbols without a tilde represent concentration.

In this study, we establish a connection between experimental observations and the MCRD model to elucidate the mechanisms of phase separation. Our theoretical predictions, validated through experiments, reveal that mesoscale biomolecular condensates exhibit local concentration-dependent feedback. By incorporating experimentally derived parameters, we develop an MCRD model that accurately reconstructs key features of DPIC formation, including plateau-shaped concentration profiles, phase diagrams, and the spatiotemporal scale-invariance of DPICs, all in alignment with experimental data. Notably, the MCRD model predicts trends in the exponential factor governing the growth of DPIC formation, which agree with experimental observation. Dispersion relation analyses, derived from both experimental data and model simulations, confirm that DPICs undergo coarsening during the early stages of development and exhibit instability of the homogenous state. This model also demonstrates that the DPICs exhibit self-similar dynamical properties at the latest time available. These insights offer new perspectives on MCRD systems in modelling phase separation. Using this simplified mechanistic model, we may uncover curious processes that drive the formation of biomolecular condensates in living cells.

## Results

### DPICs exhibit local concentration-dependent dynamics of proteins

Based on DPICs experiment data (Section 1.1 in **Supplementary Methods**) and the previous work (*47*), dsDNA primarily functions as a scaffold that promotes the conversion of free proteins into dsDNA-bound proteins. The role of dsDNA is treated as a parameter in our dynamic model of DPICs. Regions enriched with dsDNA-bound proteins represent the DPICs, while areas with low densities of dsDNA-bound proteins are considered outside the DPICs. We introduced two variables defined directly as the concentrations of dsDNA-bound protein (*u*) and free protein (*v*) (Section 1.1 in **Supplementary Methods**) to capture the dynamic behavior of DPICs formation at meso-scale. Our goal is to model the dynamics or response function governing the exchange between them (Fig. 1A(ii)). It is pertinent to note that in the subsequent sections, mathematical symbols with a tilde denote the fluorescent intensity measured from experiments, whereas symbols without a tilde represent concentration. A strong positive correlation between the average intensity of fluorescent molecules and their concentration is established in Supplementary Fig. 4A to guarantee the conversion between them.

To directly test the response functions of local concentration-dependent behavior within DPICs, where dsDNA-bound proteins recruit additional free proteins, we devised an *in vitro* droplet assay, termed the “recruitment experiment” (Supplementary Fig. 3A). Initially, we mixed 20 μM unlabeled p53^4M^ ΔTAD with 0.6 μM FAM-labeled 400-bp random dsDNA for a 20-minute incubation to generate DPICs. Subsequently, we removed all unbound protein and dsDNA. Finally, we introduced 20 μM ATTO565-labeled p53^4M^ ΔTAD and observed a rapid infiltration of red-fluorescent proteins into the original green DPICs (Fig. 1B(i)). The time traces of ATTO565-labeled p53^4M^ ΔTAD intensities were quantified, revealing that protein recruitment reached quasi-equilibrium after approximately one hour (N = 41, Supplementary Fig. 3B). We found a significant positive correlation (Fig. 1B(ii)) between the intensity of recruited ATTO565-labeled p53^4M^ ΔTAD at equilibrium (3,540 s in Supplementary Fig. 3C) and the initial amount of p53^4M^ ΔTAD inside the DPIC (0 s, Supplementary Fig. 4B(v) and **Methods**). This indicates a positive feedback mechanism between protein recruitment and local concentration of protein. A recruitment function, 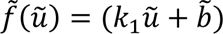, was used to fit the experimental data well: (i) *k*_1_ = 1.35 ± 0.05, the recruitment response coefficient per increment unit *u* (Supplementary Table 1); (ii) *b* = 93 ± 56, the self-triggered rate when absence of dsDNA-bound proteins (*u* = 0) (Supplementary Table 1). So that 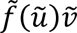 express the nonlinear recruitment mechanism.

To quantify the presence of local concentration-dependent negative feedback within DPICs, where protein escape is influenced by the stability of dsDNA-bound complexes and local concentration gradients, we devised a series of *in vitro* droplet assays termed “escape experiments” (Supplementary Fig. 5A). Initially, we incubated ATTO565-labeled p53^4M^ ΔTAD and FAM-labeled 400-bp random dsDNA for 45 minutes at room temperature to generate large dual-color DPICs. After removing all unbound protein and dsDNA, we introduced a blank working buffer and began recording the time traces of ATTO565-labeled p53^4M^ ΔTAD intensities. We prepared a total of 15 different experimental conditions of DPICs by varying the initial protein and dsDNA concentrations (Supplementary Fig. 5B), selecting the condition with 10 μM ATTO565-labeled p53^4M^ ΔTAD and 0.3 μM FAM-labeled 400-bp random dsDNA as a representative example (Fig. 1C(i)). We observed a gradual decrease in protein intensity under this condition, consistent with all other experimental setups (Supplementary Fig. 5C). To mitigate potential photobleaching effects, we collected data at only six time points from 0 to 18 hours post-injection. The declining protein intensity suggested that protein molecules were indeed escaping from the DPICs. This escaping behavior stabilized after 18 hours (Supplementary Fig. 5D), with the escape ratio of proteins at the 18-hour time point showing a pronounced negative correlation with the normalized initial amount of p53^4M^ ΔTAD inside DPIC at 0-hour time point (Fig. 1C(ii)). This indicates a negative feedback mechanism between the ratio of protein escaped from the DPIC and the local concentration of protein inside the DPIC. An escape function, 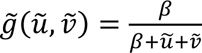, was used to fit these experimental data well: *v* = 0.5 (A small value from the experimental estimation), *β* = 1.6 ± 0.4, the half-saturation constant (Supplementary Table 1).

The detachment process can be quantified by measuring the persistence time distribution of dsDNA-bound proteins. This approach provides a direct experimental method for estimating the detachment rate constant, *k*_*d*_, from observed DPIC. This parameter has been measured by experiment (*49*), demonstrating that the detachment kinetics can be accurately characterized by analyzing the temporal behavior of individual molecules bound to dsDNA.

Taken together, our p53-dsDNA DPIC system encompasses two types of response functions: the first describes the participation of free proteins in the formation of DPICs, while the second delineates the transformation of DPIC-bound proteins back into free proteins, as well as the spontaneous detachment process within nonlinear biochemical reactions. We have tested different nonlinear rates for *f*(*u*) and *g*(*u*, *v*) using AIC and BIC criteria (Supplementary Table S2); however, they did not result in significant differences in our findings.

### A mass-conserving reaction-diffusion model for DPICs

To understand the formation behaviors of DPICs and establish a mass-conserving dynamical model based on our experiments, we utilized the two response functions discussed earlier (Fig. 2A(i)). For any given spatial point (*x*, *y*) at time *t*, the change of dsDNA-bound protein, *u*(*x*, *y*, *t*), depend on concentration of components within radius *Δx* (Fig. 2A(ii)), and the profiles of *u*(*x*, *y*, *t*) reflects where the proteins are clustered to form DPICs. The dynamics of *u* and *v* are governed by the equation (1) (Fig. 2A(iii)):

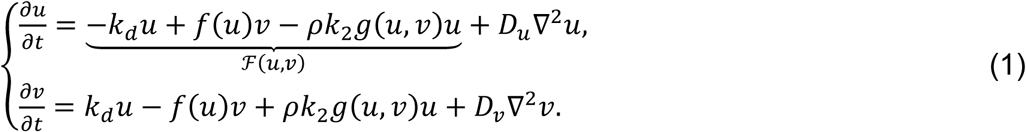

**Fig. 2.**
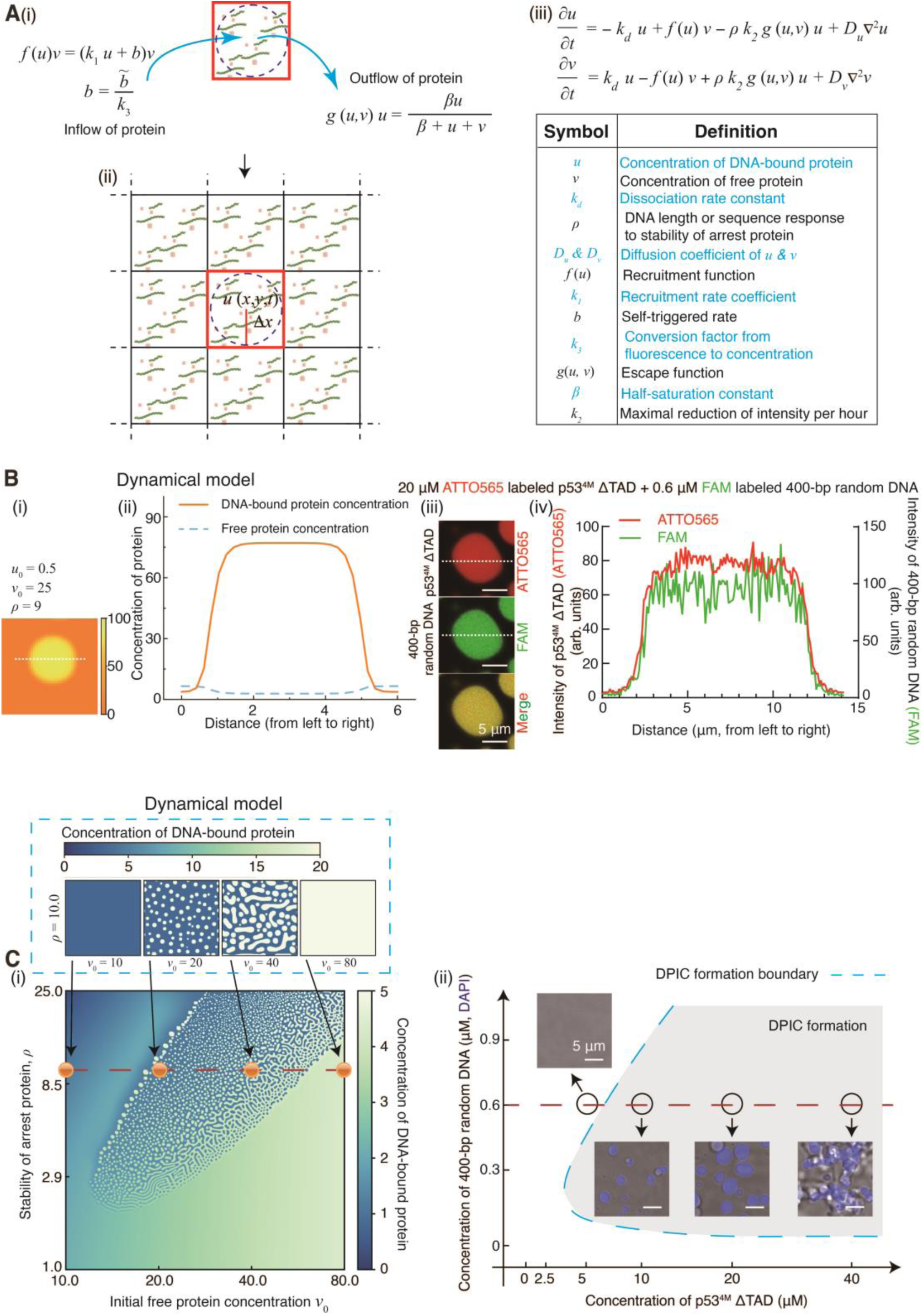
A mass-conserving dynamical model successfully reproduced the spatiotemporal dynamics of DPICs. (**A**) A mass-conserving dynamical model was established. (i) A diagram illustrating the kinetic processes of recruitment and escape at space (*x*, *y*) within a Δ*t* time interval; (ii) Each spatial lattice point is associated with the protein-dsDNA interactions that are analogous to a spatially homogeneous system; (iii) The mass-conserving dynamical model. All symbol definitions were in the table (Supplementary Table 1 and **Supplementary Methods**). (**B**) (i) A simulated droplet-like DPIC by a numerical simulation with *u*_0_ = 0.5, *v*_0_ = 25, *ρ* = 0.5, *D*_*v*_ = 0.17, *D*_*u*_ = 0.05*D*_*v*_; (ii) The simulated DPIC profiles for dsDNA-bound protein (*u*) and free protein (*v*) in (i) showed a plateau shape; (iii) A experimental droplet-like DPIC by a two-color *in vitro* droplet assay involving the mixing of 0.6 μM 400-bp random dsDNA labeled by FAM with 20 μM p53^4M^ ΔTAD labeled by ATTO565; (iv) The experimental phase-separation profile in (iii) also showed a plateau shape. (**C**) (i) Model-predicted phase diagram, which was plotted in the plane of *v*_0_ and *ρ*. (*Insets*) Model-predicted spatial patterns following a red dashed line by fixing the value of *ρ* = 10 and varying *v*_0_(10, 20, 40, and 80); (ii) Experimental phase diagram, which was plotted in the plane of concentrations of p53^4M^ ΔTAD and concentration of 400-bp random dsDNA. (*Insets*) Experimental spatial patterns following a red dashed line by fixing the value of 0.6 μM 400-bp random dsDNA and varying p53^4M^ ΔTAD concentrations (5, 10, 20, and 40 μM).

The total mass of proteins is conserved, i.e.,*τ* = ∫_Ω_ [*u*(*x*, *t*) + *v*(*x*, *t*)]*d****r*** is constant (Ω is the confined space). The reaction kinetics ℱ(*u*, *v*) comprise detachment, and escape processes of dsDNA-bounded proteins, and recruitment of free proteins by DPICs (Section 1.1 in **Supplementary Methods**).

Here, we described each term in Eq. 1:

(i) The 1^st^ term represents that protein dissociates from dsDNA, which can be written as *k*_*d*_*u*, where *k*_*d*_ is the dissociation rate constant. We assume the *k*_*d*_ value of p53^4M^ ΔTAD-dsDNA is equal to that of wild type p53-dsDNA, which was reported as 0.40 ± 0.13 *s*^−1^ in the reference (Supplementary Table 1) (*49*).

(ii) The 2^nd^ term is the recruitment of free protein by the DPIC. This item is expressed as nonlear function of *f*(*u*)*v*, in which *f*(*u*) is the recruit function (Figs. 2A(i) & 1B). *b* = *b*⁄*k*_3_, and *k*_3_ is conversion factor from fluorescence to concentration (Supplementary Fig. 4B(vi) and **Methods**).

(iii) The 3^rd^ term is the escape of the dsDNA-bound protein from the local area to the surrounding buffer. Considering that the concentration, length, and sequence of dsDNA can change the structure of DPIC and affect the movement of dsDNA-bound protein, we used the parameter *ρ* to capture the stability effects of dsDNA-bound protein properties here. Therefore, this item is written as *ρg*(*u*, *v*)*u*, where *g*(*u*, *v*) is the escape function (Figs. 2A(i) & 1C), and *k*_2_ is maximal reduction of intensity per hour (Estimated from Supplementary Fig. 5D).

(iv) The 4^th^ term describes the Brownian movement of species, and ∇^2^ is the Laplacian operator, i.e., 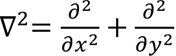. Our FCS experiments provide *D*_v_ = 18.0 ± 1.1 *μm*^2^⁄*s*, which matches the value of *D*_*v*_ = 15.4 ± 5.6 *μm*^2^⁄*s* provided in the previous study (Supplementary Table 1) (*49*). Our previous FRAP experiments (*47*) provide *D*_*u*_ = (9.2 ± 7.4) · 10^−4^ *μm*^2^⁄*s* (Supplementary Fig. 2C), which is much smaller than *D*_*v*_.

In the above four terms, the positive recruitment led to the local increase of *u*, while the protein dissociation and escape lead to the decrease of *u*. The nonlinear functions of *f*(*u*) and *g*(*u*, *v*) lead to the broken detailed balance at the mesoscopic scale. Due to the conservation of reactions, the contribution of the first three terms to the change of *v* is exactly the opposite of their contribution to that of *u*.

### Dynamical model reproduces the spatiotemporal dynamics of DPICs

**(i) The plateau-shaped concentration profiles and phase diagram of DPICs.** We initiated the simulation with *v*_0_ = 25 and *ρ* = 9 (**Methods** and Table 1). Upon reaching equilibrium, the dsDNA-bound proteins formed condensates (Fig. 2B(i)), with a distinct concentration boundary separating the inside and outside the DPIC (Fig. 2B(ii)). The simulation revealed a plateau-shaped concentration profile inside the condensates, closely matching the results from the droplet-like DPIC experiments (Fig. 2B(iii)-(iv)).

**Table 1.**
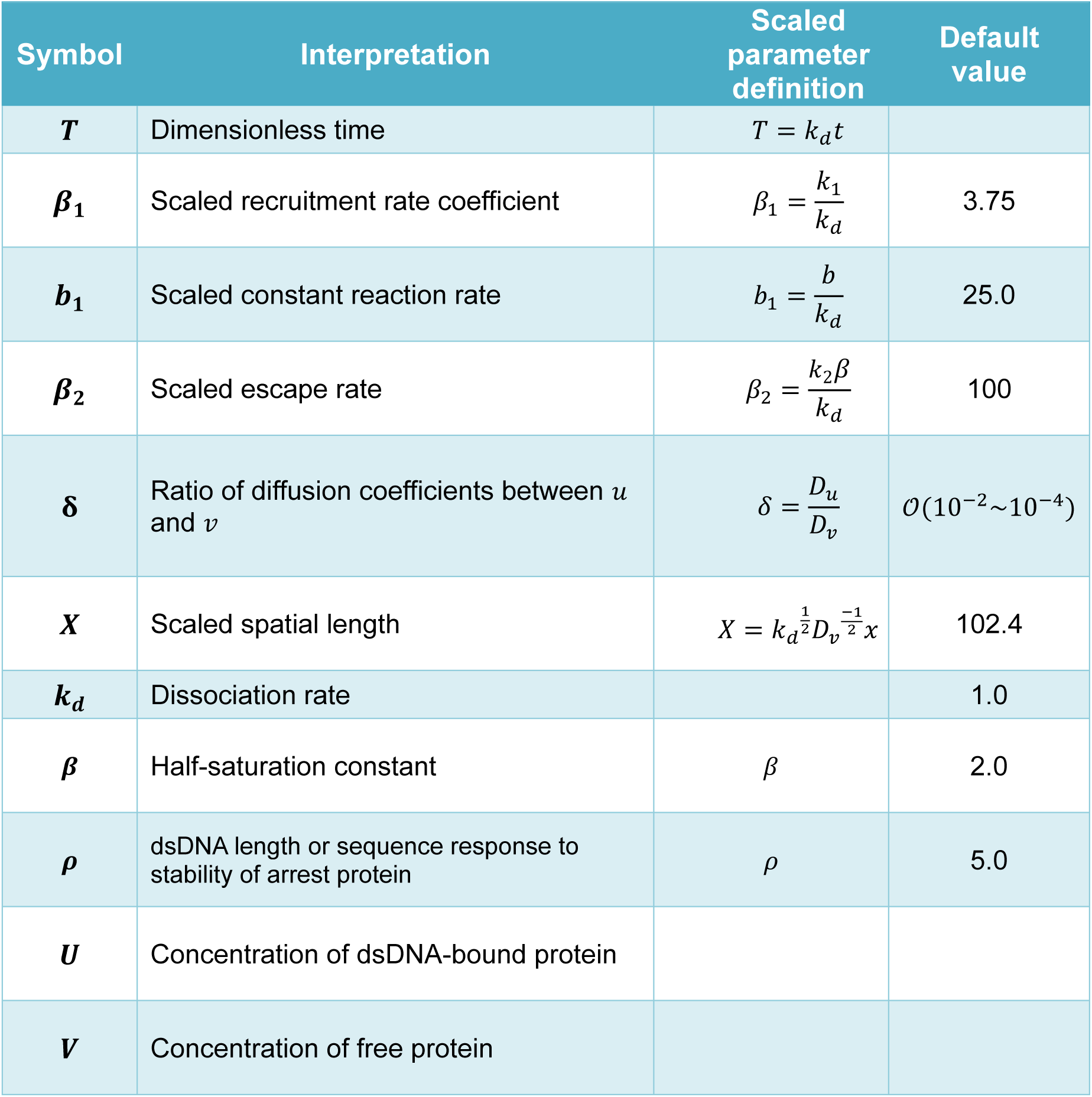
Scaled parameter interpretations, default values, and their definitions.

To further investigate, we set the initial values of *v*_0_ and *ρ* as parameter gradients along the transverse and longitudinal axes respectively (Section 1.2 in **Supplementary Methods**). This allowed us to construct a phase diagram (Fig. 2C(i)). By fixing the value of *ρ* = 10 and varying *v*_0_ (10, 20, 40, and 80, as indicated by the red dashed line in Fig. 2C(i)), we observed different pattern formations (Fig. 2C(i) and insets). At a low initial *v*_0_ (10), no condensates were formed. As *v*_0_ increased to 20 and 40, droplet-like and “pearl chain”-like condensates appeared, respectively. When *v*_0_ was further increased to 80, condensates were no longer formed again. Our experimental phase diagram (Fig. 2C(ii) and Supplementary Fig. 1D) exhibited similar properties. We mixed 0.6 μM of 400-bp random dsDNA with varying concentrations of p53^4M^ ΔTAD. At a low protein concentration (5 μM), no DPICs were formed. When the protein concentration increased to 10 μM and 20 μM, droplet-like DPICs were observed. Further increasing the protein concentration to 40 μM resulted in the formation of “pearl chain”-like DPICs.

**(ii) Spatiotemporal scale-invariance of DPICs.** To further understand the spatiotemporal self-organized patterns of DPICs, we analyzed the temporal evolution of the spatial wavelength of DPIC patterns from our experiments (Figs. 3A, D, and Session 1.3 in **Supplementary Methods**). This analysis provides two key parameters: (i) Spatial scales ℓ, characterized by the time-dependent average wavelength ℓ = 2*πL*/*q*_*max*_ (where *L* represents the window size, determined using a moving window method); (ii) Dominant wavenumbers *q*_*max*_(*t*) = ∫ *qS*(*q*, *t*) *dq*⁄∫ *S*(*q*, *t*)*dq*, where *q* represents the wavenumber and *S*(*q*, *t*) is the dynamic structure factor (Section 1.3 in **Supplementary Methods**) (*21, 50*).

**Fig. 3.**
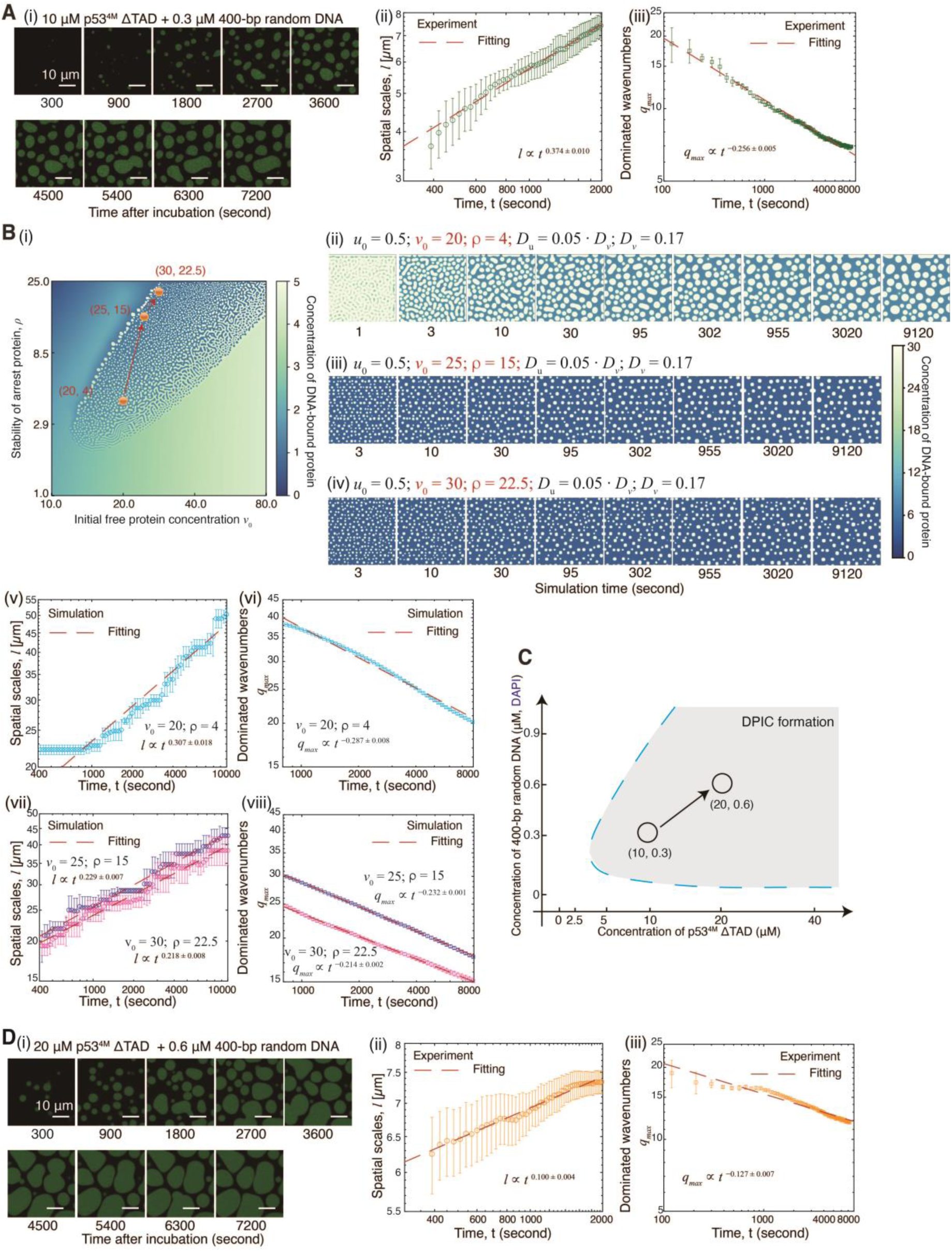
Spatiotemporal scale-invariance of DPICs from experiments and numerical simulations. (**A and D**) Spatiotemporal dynamics of DPICs formed by 0.3 μM 400-bp random dsDNA labeled by FAM with 10 μM dark p53^4M^ ΔTAD (A), or 0.6 μM 400-bp random dsDNA labeled by FAM with 20 μM dark p53^4M^ ΔTAD (D). (i) Time course of coarsening behavior of DPICs, with time indicated in seconds; (ii) Spatial scales ℓ over time; (iii) Dominant wavenumbers *q*_*max*_ over time. Independent analysis was repeated: N = 5 for 5 different observation regions in one experiment. Both the spatial scales and dominant wavenumbers follow power-law relations (The dashed line in (ii)-(iii) represents the power-law fitting). Error bars, mean ± s.d. (**B**) Spatiotemporal dynamics of simulated DPICs. (i) Model-predicted phase diagram, which was plotted in the plane of *v*_0_ and *ρ*. (ii)-(iv) Simulation time course of coarsening behavior of DPICs for three different model conditions, with simulation time indicated in seconds: (ii) *u*_0_ = 0.5, *v*_0_ = 20, *ρ* = 4.0, *D*_*v*_ = 0.17, *D*_*u*_ = 0.05*D*_*v*_; (iii) *u*_0_ = 0.5, *v*_0_ = 25, *ρ* = 15.0, *D*_*v*_ = 0.17, *D*_*u*_ = 0.05*D*_*v*_; (iv) *u*_0_ = 0.5, *v*_0_ = 30, *ρ* = 22.5, *D*_*v*_ = 0.17, *D*_*u*_ = 0.05*D*_*v*_. The color bar represents the concentration of dsDNA-bounded protein. The direction of the pathway for these three different model conditions is upper right. (v)-(viii) Spatial scales ℓ and dominant wavenumbers *q*_*max*_ over time. (v)-(vi) for model condition (ii); (vii)-(viii) for model condition (iii)-(iv). Independent simulations were repeated 5 times (n = 5) for (ii)-(iv). Error bars, mean ± s.d. (**C**) Experimental phase diagram (Fig. 2C(ii) and Supplementary Fig. 1D). The direction of the pathway for these two different experimental conditions is also upper right.

We mixed 0.3 μM of 400-bp random dsDNA with 10 μM p53^4M^ ΔTAD and observed the dynamical behaviors over a two-hour period (Fig. 3A(i) and Supplementary Movie 1). The experimental video demonstrated that both spatial scales and dominant wavenumbers (Section 1.3 in **Supplementary Methods**) are time-dependent. Interestingly, the spatial scales follow a power-law relation scale ℓ ∝ *t*^*m*^, and the dominant wavenumbers follow *q*_*max*_ ∝ *t*^−*n*^ (Fig. 3A(ii)-(iii)). These results indicate that DPICs exhibit spatiotemporal scale-invariance. Specifically, the exponential factor *m* for the spatial scales was approximately 0.37, while for the dominant wavenumbers, *n* was approximately 0.26 (Fig. 3A(ii)-(iii)). To further validate our findings, we selected a model condition with *v*_0_ = 20 and *ρ* = 4 from the phase diagram (Fig. 3B(i)) and performed numerical simulations of the coarsening behaviors (Fig. 3B(ii) and Supplementary Movie 2). The simulation results showed an exponential factor *m* = 0.31 for the spatial scales and *n* = 0.29 for the dominant wavenumbers (Fig. 3B(v)-(vi)). These results demonstrate that our dynamical model successfully reproduces the spatiotemporal scaling dynamics of DPICs. For patterns arising from a Turing instability, their spatial scales and dominant wavenumbers become time-independent at long times. However, for patterns exhibiting coarsening behaviors, both the spatial scales and dominant wavenumbers are always time-dependent, typically with a power-law in time at long enough times. This implies that DPICs exhibit a dynamic spatiotemporal scaling behavior with coarsening.

To investigate whether we can predict the trend of the exponential factor of the dominant wavenumbers under different model conditions, we selected a pathway in the phase diagram. This pathway transitions from an initial condition of *v*_0_ = 20 and *ρ* = 4 to *v*_0_ = 25 and *ρ* = 15, and finally to *v*_0_ = 30 and *ρ* = 22.5 (Fig. 3B(i)). The coarsening behaviors along this pathway were simulated (Fig. 3B(iii)-(iv) and Supplementary Movies 3-4). We observed that the exponential factor *m* of the spatial scales decreased from 0.31 to 0.23, and then to 0.22, and the exponential factor *n* of the dominant wavenumbers decreased from 0.29 to 0.23, and then to 0.21 (Fig. 3B(vii)-(viii)). To confirm this prediction experimentally, we chose a pathway with a similar direction in the phase diagram (Fig. 3C and Supplementary Fig. 1D). Starting from the condition of 0.3 μM 400-bp random dsDNA with 10 μM p53^4M^ ΔTAD, we moved to 0.6 μM 400-bp random dsDNA with 20 μM p53^4M^ ΔTAD (Fig. 3D(i) and Supplementary Movie 5). the exponential factor *m* of the spatial scales decreased from 0.37 to 0.10, and the exponential factor *n* of the dominant wavenumbers decreased from 0.26 to 0.13 (Fig. 3D(ii)-(iii)), which aligns well with the predictions from our dynamical model.

Taken together, our dynamical model successfully reproduces the plateau-shaped concentration profiles, phase diagram, and spatiotemporal scale invariance of DPICs. These results demonstrate that mass-conserving dynamical models, previously applied to understanding animal and pebble aggregations, can also elucidate the mechanisms underlying DPIC formation. Thus, both mesoscale biomolecular condensates and macroscale biogeomorphological systems can be understood through a unified MCRD model.

### Dispersion relation reveals coarsening behavior of DPICs in the early stage

We performed a two-dimensional (2D) spectral decomposition of our experimental DPIC patterns (Fig. 3A(i) and Supplementary Movie 1) to isolate the contribution of individual wavelengths (or wavenumbers) to the overall coarsening patterns (Section 1.5 in **Supplementary Methods**). We selected three specific wavelengths—*λ* = 578.0 μm, 60.9 μm, and 17.0 μm—as examples, where *λ* = 2*π*/*q*. By focusing on a time window from 300 seconds to 700 seconds, which corresponds to the early stage of the 7,200-second experimental video (Supplementary Movie 1), we observed that the calculated amplitude (Section 1.5 in **Supplementary Methods**) varies exponentially with time (Fig. 4A(i)-(iii)). The slope of the logarithmic amplitude versus time indicates the growth rate σ(*q*) of the selected wavenumber *q*. We then plotted the growth rate σ(*q*) as a function of each wavenumber *q*, obtaining the dispersion relation (the blue data points in Fig. 4B). Notably, the experimental dispersion relation started from a positive value when the value of *q* was small.

**Fig. 4.**
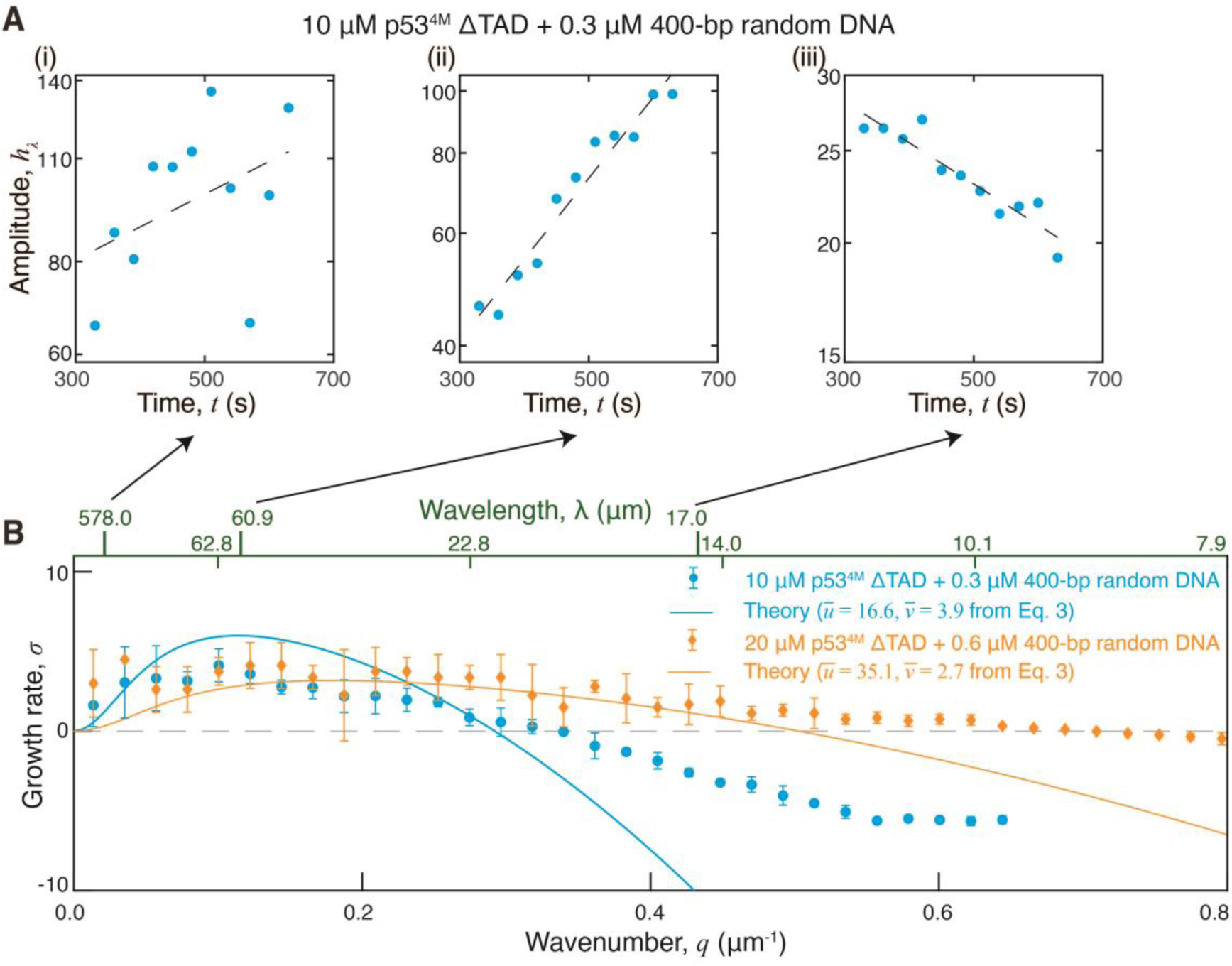
The dispersion relation of DPIC from experiments and theoretical predictions. (**A**) For the DPICs of 0.3 μM 400-bp random dsDNA with 10 μM p53^4M^ ΔTAD, amplitude versus time (300-700 seconds, the early stage of the 7,200-second experimental video) for three specific wavelengths— *λ* = 578.0 μm (i), 60.9 μm (ii), and 17.0 μm (iii)—as examples, where *λ* = 2*π*/*q*. The slope of the logarithmic amplitude varies exponentially with time (the black dashed line in (i)-(iii)) indicates the growth rate σ(*q*) of the selected wavenumber *q*. (**B**) The growth rate σ(*q*) versus wavenumber *q* (and wavelength *λ*). Experimental condition 1 (blue circle): 0.3 μM 400-bp random dsDNA with 10 μM p53^4M^ ΔTAD; Model condition 1 (blue line): *u*̅ = 16.6, *v*̅ = 3.9, *ρ* = 4.0, *δ* = 0.003; Experimental condition 2 (orange circle): 0.6 μM 400-bp random dsDNA with 20 μM dark p53^4M^ ΔTAD; Model condition 2 (orange line): *u*̅ = 35.1, *v*̅ = 2.7, *ρ* = 16.0, *δ* = 0.0005. The direction of the pathway from Experimental condition 1 to Experimental condition 2 in the phase diagram is the same as from Model condition 1 to Model condition 2. Error bars, mean ± s.d.

We next asked whether our established mass-conserving dynamical model can theoretically predict our experimental dispersion relation. For this purpose, we interrogate the stability of the homogeneous steady state of Eq. 4 (**Methods**) with respect to wave-like perturbations through standard linear stability analysis, by linearizing Eq. 4 around this steady state. Every perturbation can be represented as a linear combination of Fourier modes, which initially evolve independently from each other. Linearizing and taking a Fourier transform 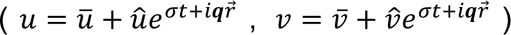 (Section 1.4 in **Supplementary Methods**), we obtain the characteristic equation

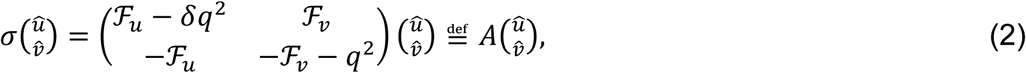

where σ is temporal frequency and ***q*** = (*q*_*x*_, *q*_*y*_) is the wave vector, which can also be expressed in terms of the perturbation wavelength *λ* = 2*π*/*q* with 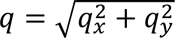. Note that the perturbations must respect the mass constraint of system Eq. 4. Self-organized patterns occur for wavenumber *q* > 0 with positive real part of σ(*q*). In this case, as expected, the most unstable perturbations are the fixed undulations at the scale of *q* = 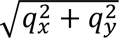 with the maximum growth rate ℜ*e*[σ(*q*)], where

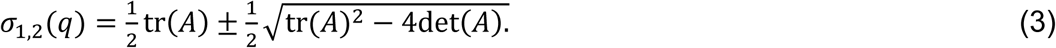

Eq. 3 provides the theoretical dispersion relation for the model described by Eq. 4, illustrating the initial rate of growth or decay of perturbations with various wavenumbers (Section 1.6 in **Supplementary Methods**). From Fig. 3, we determined that the experimental condition (0.3 μM 400-bp random dsDNA with 10 μM p53^4M^ ΔTAD) corresponds to a model condition with *u*_0_ = 0.5, *v*_0_ = 20 and *ρ* = 4 in the phase diagram (Fig. 3B(i)). Using these data, we plotted the theoretical dispersion relation ℜ*e*[σ(*q*)] as a function of wavenumber *q* based on Eq. 3 with *u*̅ = 16.6 and *v*̅ = 3.9 (the solid line in Fig. 4B), which aligns well with the experimental data (the data points in Fig. 4B).

To investigate whether we could predict the trend of the dispersion relation, we adjusted the model condition to *u*_0_ = 0.5, *v*_0_ = 37.3 and *ρ* = 16 in the phase diagram (Fig. 3B(i)). The theoretical prediction for this condition (represented by the brown solid line) is plotted in Fig. 4B based on Eq. 3 with *u*̅ = 35.1 and *v*̅ = 2.7. Considering the total protein concentration (*u*̅ + *v*̅) was increased from 20.5 to 37.8, to experimentally confirm this prediction, we selected an experimental condition of 0.6 μM 400-bp random dsDNA with 20 μM p53^4M^ ΔTAD (Fig. 3D(i) and Supplementary Movie 5), which corresponds to a similar direction of increased total mass of protein with the model condition in the phase diagram (Fig. 3C). Comparing the theoretical dispersion relation with the experimental data (represented by the orange dots in Fig. 4B), we observed that the experimental data exhibited the same characteristic shape as the theoretical prediction.

Taken together, our findings reveal that DPICs exhibit a dynamical pattern through the specific dispersion relation. Furthermore, our analysis demonstrated that DPICs form spatiotemporal self-organized patterns with coarsening behaviors during the early stages. These conclusions collectively provide compelling evidence that dispersion relation reveals coarsening behavior of DPICs in the early stage.

### DPICs exhibit self-similar dynamical properties

When we analyzed the dominant wavenumbers *q*_*max*_ (Section 1.3 in **Supplementary Methods**) for the phase separation system of 0.3 μM 400-bp random dsDNA and 10 μM p53^4M^ ΔTAD (Fig. 3A(i) and Supplementary Movie 1), we discovered a notable result. By plotting *S*(*q*, *t*)*q*^2^_*max*_, defined as the structure function, versus rescaled *q*⁄*q*_*max*_ at various time points, we observed that each structure function follows a power-law relation. Remarkably, all data points collapsed onto a single universal master line, with an exponential factor of -4.0 (Fig. 5A(i)). Furthermore, a different experimental condition of 0.6 μM 400-bp random dsDNA and 20 μM p53^4M^ ΔTAD (Fig. 3D(i) and Supplementary Movie 5) has the same exponential factor of -4.0 (Fig. 5B(i)). When we repeated this analysis for two different numerical simulations in the phase diagram (Fig. 3B(i)), *v*_0_ = 20 and *ρ* = 4; *v*_0_ = 30 and *ρ* = 22.5, we obtained the same exponential factor of -4.0 (Fig. 5A(ii) and B(ii)). These results suggest that at the latest times available, the form factor approaches a Porod’s tail form for large wavenumber (*50*).

**Fig. 5.**
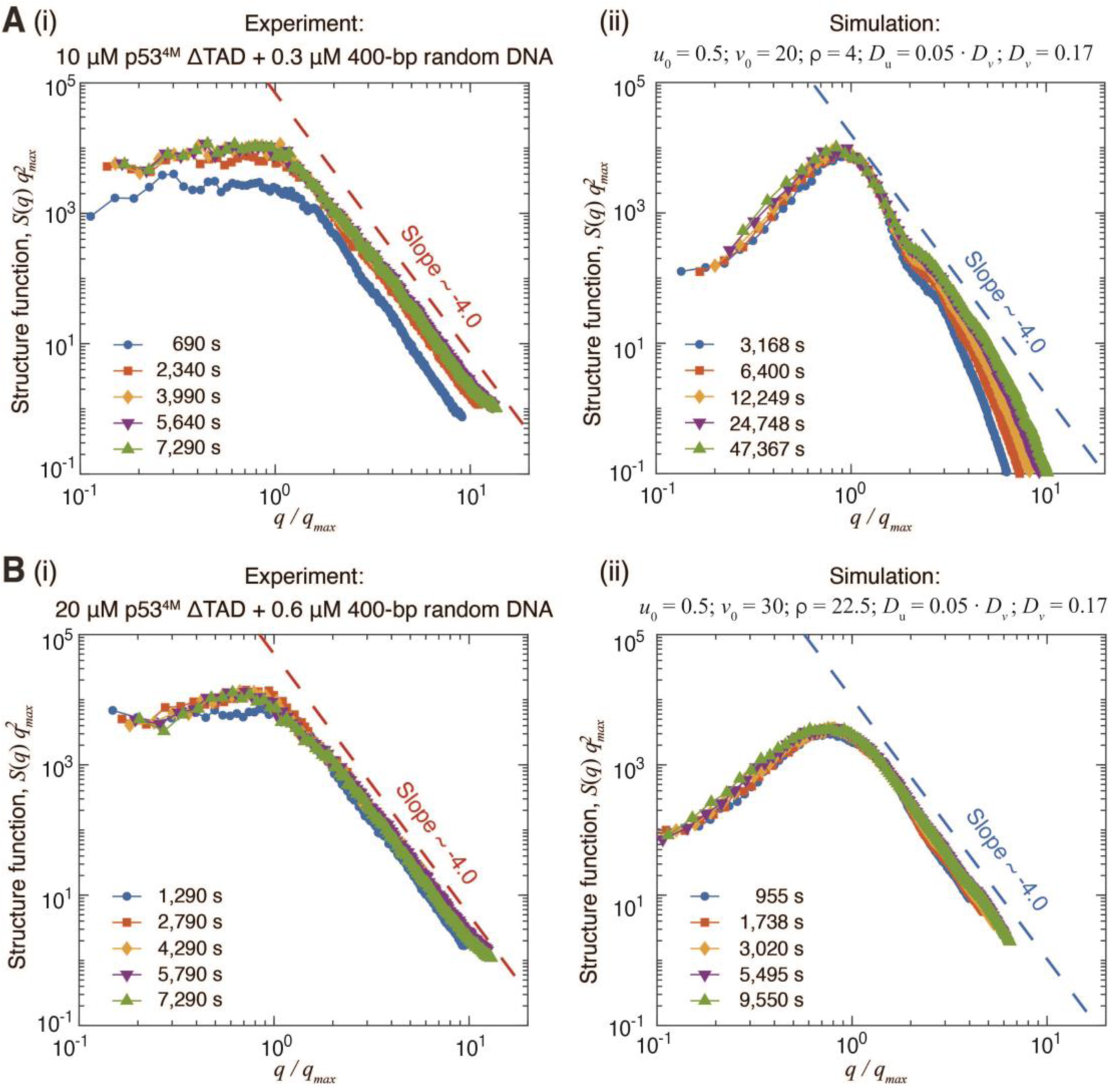
Self-similar dynamical properties of DPICs from experiments and numerical simulations. (**A-B**) (i) The experimental data were plotted as the rescaled structure unction *S*(*q*)*q*^2^_*max*_ versus *q*/*q*_*max*_ at various time points. All points collapse onto one single power-law line (dashed red lines) with a same slope ∼-4.0. (ii) We used two different model conditions to conduct numerical simulations, reproduced the analysis in panels A(i) and B(i). The rescaled structure factors collapse to form a master line all with the same slope ∼-4.0 (N = 10 for the mean values). 10 μM p53^4M^ ΔTAD mixed with 0.3 μM 400-bp random dsDNA (A); 20 μM p53^4M^ ΔTAD mixed with 0.6 μM 400-bp random dsDNA (B).

## Discussion

In this study, we conducted a quantitative analysis of a specific type of mesoscale biomolecular condensate, known as DPICs, by establishing an MCRD model based on our experiments. Our findings demonstrate that DPICs exhibit plateau-shaped concentration profiles (in contrast to peak patterns observed in biological pattern formation (*51*)) and phase diagrams (Fig. 2), spatiotemporal self-organized patterns with coarsening behaviors during the early stages (Figs. 3-4), and self-similar dynamical properties (Fig. 5). Through the mesoscale experimental feedback, we developed an MCRD model which can reproduce the phase separation patterns observed in *in vitro* DPIC experimental system. Furthermore, the MCRD model can also qualitatively predict the dynamic behavior of DPIC—the increased stability of protein on dsDNA (*ρ*) and the increased concentration of protein (*v*_0_) could slow down the growth rate of droplets (Fig. 3). Consequently, DPIC system bridges the experimental and theoretical studies of MCRD systems in phase separation. Based on our work, several interesting topics warrant further exploration in future studies.

First, both our experimental evidence and theoretical modeling indicate that MCRD-type equations exhibit a general coarsening behavior, a hallmark of phase separation consistently observed across experimental and theoretical studies (*21, 52*). These findings suggest that MCRD models offer a promising alternative framework for exploring phase separation in diverse systems. Notably, this coarsening behavior persists under approximately mass-conserving conditions, as demonstrated in recent studies on cell polarity aggregation (*53*). The robustness of MCRD models under such conditions underscores their potential as a valuable paradigm for investigating pattern formation in mass-conserved systems exhibiting coarsening dynamics. Further quantitative investigation from both experimental and mathematical perspectives is warranted to expand this area of research.

Second, it is worth emphasizing that the MCRD model has been widely applied to active biological and chemical systems to uncover the mechanisms underlying pattern formation (*10, 38, 54, 55*). In this study, we extend the MCRD framework to describe biomolecular condensates, focusing on reversible autocatalytic biochemical reactions. Unlike prior approaches that integrate thermodynamic processes with reaction kinetics (*27, 33, 34*), our analysis relies solely on reaction-diffusion dynamics to elucidate the coarsening behavior associated with phase separation. The DPIC system does not require ATP hydrolysis or other external energy sources. This suggests that systems composed of diverse interacting components, when placed in environments where certain configurations persist better than others, inherently evolve toward states of increasing “functional information” (*56, 57*). This insight may have implications for understanding the origins of organic matter.

Third, in Fig. 3, the spatial scales of DPICs follow a power-law relation, ℓ ∝ *t*^*m*^, demonstrating spatiotemporal scale-invariance. Remarkably, each well-studied system of animal and pebble aggregations, characterized by a unique coarsening exponent *m*, can be associated with a biomolecular condensate system exhibiting a similar coarsening exponent. For instance, both macroscale mussel beds (*20*) and mesoscale nucleolar protein FIB-1 condensates in living cells (*58*) share a coarsening exponent of 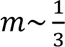 macroscale stone patterns (*19*) and mesoscale DPICs in our experiments (*47*) both exhibit 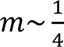; and both macroscale dunes and ripples (*59*)) and mesoscale chromatin-protein condensates in living cells (*60-62*) have coarsening exponents 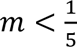. As we all know, both Ostwald ripening (*63, 64*) and Brownian motion coalescence (*65*) can promote a coarsening behavior with a coarsening exponent of 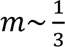. This high degree of consistency suggests a compelling idea: mesoscale biomolecular condensates could serve as model systems for studying macroscale biogeomorphological systems, akin to how *Drosophila melanogaster* (fruit fly) and *Caenorhabditis elegans* (nematode worm) are used to model human diseases. This is an interesting topic for the future study.

In summary, we establish a connection between experimental observations and theoretical modeling of MCRD systems in phase separation using a specific type of biomolecular condensate. Our findings demonstrate that MCRD systems provide a robust framework for understanding phase separation characterized by coarsening behavior. These insights offer a promising foundation for future investigations into the mechanisms driving phase separation in complex biological systems using multicomponent dynamical models. Moreover, advancing the theoretical understanding of MCRD models has practical implications for experimental biology. By integrating theory with experimentation, researchers can design and test hypotheses more effectively, leveraging an optimal combination of computational modeling and empirical validation to unravel the complexities of biological systems.

## Acknowledgments

We thank the great comments from Dr. Nigel Goldenfeld (University of California San Diego, USA). We thank Dr. Chunlai Chen (Tsinghua University, China) for help us to conduct FCS experiments. We thank the Peking Nanofab for process support. We thank the contributions of the Engineering Research Center for Semiconductor Integrated Technology, Institute of Semiconductors, Chinese Academy of Sciences. We thank all members of the Qi laboratories for comments on the manuscript.

## Author Contributions

C.L. prepared biological samples, conducted all experiments, performed data analysis, and wrote the manuscript. M.T.G. and X.Q.H. performed simulation and theoretical analysis, and X.Q.H., and Q.X.L. built up the theoretical model. Z.Q., Q.X.L., and X.Q.H. supervised the project, experimental designs, and data analysis, and wrote the manuscript with input from all authors.

## Data and code availability

All data needed to evaluate the conclusions in the paper are present in the paper and/or the Supplementary Materials.

## Materials and Methods

### Construction of bacterial expression plasmids, protein purification and protein labeling

It has been proved that the transcriptional activation domain (TAD) truncated p53 which contains 4 mutations (M133L/V203A/N239Y/N268D), called p53^4M^ ΔTAD, can form DPIC with dsDNA in *in vitro* droplet assays (*47*). We cloned p53^4M^ ΔTAD on the pRSFDuet vector and inserted a 6× His-tag to construct the expression plasmid. The sequence of 6× His-p53^4M^ ΔTAD was shown in the supplementary methods.

The protein expression plasmid was transformed to *E. coli* strain BL21(DE3), and then cultured overnight on LB agar plates at 37 °C. One single clone of bacteria was cultured overnight in 10 mL LB medium at 37 °C and 220 rpm. The culture was added into a 2 L LB medium and grew to the density OD_600_ of 0.6. The bacteria were induced to express protein by the addition of 0.3 mM IPTG and 0.1 mM ZnCl_2_, and then cultured at 16 °C and 180 rpm for 18 h. The culture was centrifuged with 4,000× g. Cells were resuspended in a lysis buffer (25 mM Tris-HCl (pH 7.5), 500 mM NaCl, 5 mM imidazole, 0.25‰ β-Mercaptoethanol (β-Me), and 5% Glycerol) with 1 mM PMSF, and then sonicated on ice. The supernatant was separated and filtered after centrifugation at 18,000× rpm for 30 min. Ni-NTA was used to bind p53^4M^ ΔTAD and then washed by a washing buffer (25 mM Tris-HCl (pH 7.5), 500 mM NaCl, 20 mM imidazole, 0.25‰ β-Me, and 5% Glycerol). The proteins were eluted with an elution buffer (25 mM Tris-HCl (pH 7.5), 500 mM NaCl, 300 mM imidazole, 0.25‰ β-Me, and 5% Glycerol), and then purified by gel filtration with a Superdex 200 increase GL 10/300 (GE Healthcare, USA) with a storage buffer (20 mM Tris-HCl (pH 7.5), 300 mM NaCl, 10% Glycerol, and 40 mM DTT). After a quick-frozen by liquid nitrogen, the proteins were stored at a -80 °C environment.

In the recruitment experiment and escape experiments, the p53^4M^ ΔTAD was labeled by ATTO565 NHS ester (Sigma Cat: 72464) at 1:1.5 molar ratios in 0.1 M NaHCO_3_ for 1 hour at 4 °C. In the FCS assay, p53^4M^ ΔTAD was labeled by Sulfo Cyanine5 (Cy5) NHS ester (Lumiprobe) at 1:4 molar ratios in reaction buffer (20 mM HEPES pH 7.5, 150 mM KCl, 10% glycerol, 1 mM TCEP) and incubated for 20 min at RT. These proteins were purified by gel filtration with a Superdex 200 increase GL 10/300 with the storage buffer, and then stored at -80 °C.

### Electrophoretic Mobility Shift Assay (EMSA)

EMSA was used to test the dsDNA-binding activity of p53^4M^ ΔTAD in this paper. 30-bp dsDNA probes were generated by annealing dsDNA primers. The top strand labeled by 5’-quasar 670 and the bottom strand with no labeling were mixed at a molar ratio of 1:1.2 in an annealing buffer (40 mM Tris-HCl (pH 8.0), 50 mM NaCl, and 10 mM MgCl_2_). The mixture was heated to 95 °C for 5 min and then slowly cooled down to room temperature (RT) together with the heater. The concentration of 30-bp dsDNA probes was considered as 1 μM. The dsDNA sequences are shown in the supplementary methods.

The dsDNA probes were diluted to 0.05 μM and the p53^4M^ ΔTAD was diluted into different working concentrations by the p53 working buffer (20 mM HEPES (pH 7.9), 150 mM NaCl, 2 mM MgCl_2_, 1 mM DTT, and 0.5 mg/mL BSA). The dsDNA probes were mixed with p53^4M^ ΔTAD, incubated at RT for 30 min, and then resolved by 1% agarose gel in TBE buffer under 60 V for 90 minutes in a cold room (4 °C). The results were scanned by an Amersham Typhoon RGB system (with a 635 nm laser and Cy5 670BP30 filter).

### *In vitro* droplet assay

The FAM-labeled 400-bp random dsDNA and Cy5-labeled 400-bp random dsDNA were obtained by polymerase chain reaction (PCR) with a template of a sequence containing no p53-specific binding sites on the genome of lambda phage. In this PCR reaction, one primer carried a FAM or Cy5 tag at the 5’ terminal, and the other primer did not carry any tag. p53^4M^ ΔTAD with or without ATTO565 labeling were prepared as above. The dsDNA and protein substrates were centrifuged at 12,000× rpm for 10 min at 4 °C and then diluted to the working concentration by ddH_2_O and p53 storage buffer, respectively. All *in vitro* samples were mixed and dropped into 384-well plates (Cellvis). The dsDNA-protein mixture was incubated at RT in a p53 droplet buffer (8 mM Tris-HCl (pH 7.5), 120 mM NaCl, 4% Glycerol, and 16 mM DTT). Confocal (Leica TCS SP8) was used to observe the DPICs.

The raw image data were analyzed using Fiji (ImageJ Version: 2.0.0-rc-61/1.52n). Fluorescence images were initially converted to 8-bit format. The average intensity of ATTO565-or FAM-labeled components was then measured by analyzing either the entire area of the DPIC or a specific region within the DPIC, depending on the experimental requirements. As demonstrated in Supplementary Fig. 4A, there is a strong linear correlation between the average intensity of fluorescent molecules and their concentration. Therefore, the concentration of protein or dsDNA in the measured region can be determined by multiplying the average fluorescence intensity of ATTO565 or FAM by a conversion factor from fluorescence intensity to concentration. Since our dynamic model is two-dimensional, the total amount of the components can be determined by multiplying their concentration by the area of measured region.

For “recruitment experiment”, the ordinate is the total intensity of ATTO565 signal recruited by each DPIC at the 3,540-second time point. According to the results of Supplementary Fig. 4B, the distribution of FAM or ATTO565 of different DPICs under the same experimental condition is highly consistent, and the average intensity of ATTO565 is 0.8 times that of FAM, so the abscissa which is the initial amount of protein in DPIC is calculated by 0.8 × total FAM intensity of DPIC at 0 s. For “escape experiment”, the ordinate is the escape ratio of ATTO565 at the 18-hour time point, which is calculated by the total intensity change over 18 hours / the total intensity at 0 h. The abscissa is the rescaled total intensity of ATTO565, which is calculated by the total intensity of each condition / the minimal value.

To estimate the value of the conversion factor ***k***_**3**_ from ATTO565 intensity to the concentration of p53^4M^ ΔTAD, we conducted the following analysis. For a mass-conserved system, the concentration of the protein and its volume fraction have the following relationship:

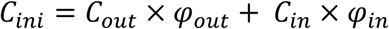

*C*_*out*_ and *C*_*in*_ represent the protein concentration outside and inside the DPIC, respectively. *φ*_*out*_ and *φ*_*in*_ represent the volume fraction of outside and inside the DPIC, respectively. *C*_*ini*_ is the initial protein concentration, which is 20 μM in the condition of Supplementary Fig. 4B. Since the volume fraction of DPIC in the whole reaction system was small, we assumed *φ*_*in*_ was 0.4%. Then, we measured *C*_*in*_/*C*_*out*_ of p53^4M^ ΔTAD by calculating the average ATTO565 intensity inside DPIC / that outside DPIC (Supplementary Fig. 4B(vi)). In this way, we estimated that the protein concentration inside DPIC was approximately 530 μM. Due to the average ATTO565 intensity inside DPIC was about 80 (Supplementary Fig. 4B(v)), the value of ***k***_**3**_ was calculated by 530 / 80, which approximated to 6.7.

### Fluorescence recovery after photobleaching (FRAP)

The ATTO565 labeled p53^4M^ ΔTAD and FAM labeled 400-bp random dsDNA were mixed and incubated for 2 hours at room temperature to obtain large DPICs. 488-nm and 561-nm lasers of a spinning-disk confocal microscope (UltraView VoX) were used to bleach an area of 1.5 μm × 1.5 μm inside DPIC with the strength of 100% for 30 cycles. The recovery process was then recorded with a speed of 2 min per frame for 120 min. A region with a radius of 1.2 μm was selected to completely cover the bleached area, which was defined as a Region Of Interest (ROI). Meanwhile, an area of the same size was selected inside DPIC without bleaching as Control. After that, the average intensity of FAM and ATTO565 in the regions of all ROIs and Controls at each time point was measured. To correct the photobleaching during the 120-minute observation, the intensity of ROI was normalized to the intensity of control. Then, the intensity of each time point of ROIs was subtracted from the intensity of the first frame after bleaching to normalize the background value to 0. Finally, the results were divided by the initial intensity before photobleaching to normalize the initial value to 1.

The FRAP result was shown in the previous work (*47*). We then re-analyzed the data to calculate the apparent diffusion coefficient (*D*_*app*_) of p53^4M^ ΔTAD inside DPIC (*66*). The normalized curves of ROIs were first fitted by *I*_(*t*)_ = *A* · (1 − *e*^−*τt*^). *I*_(*t*)_ is the normalized protein intensity of each time point, *A* is the mobile fraction of protein and *τ* is the recovery time constant. Then, the half time of fluorescent recovery *T*_1/2_ is calculated by *T*_1/2_ = 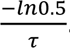. Finally, the *D*_*app*_ is calculated by 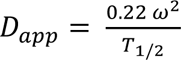 indicates the radius of the actual bleached region, which is 1.2 μm in our work.

### Fluorescence Correlation Spectroscopy (FCS)

The FCS measurement was performed by a home-made confocal microscope. The detailed information about this setup can be referred to the previous work (*67, 68*). The power of 640nm-laser after the objective was set as 5 μW. The laser focus was 10 μm above the interface of coverslip and water. Cy5-labeled p53^4M^ ΔTAD and Cy5-labeled 400-bp random dsDNA were diluted to 20 nM by an FCS buffer (8 mM Tris-HCl (pH 7.5), 120 mM NaCl, and 16 mM DTT) and then loaded into the flow cell. Raw data of photon arriving time of Cy5 detection channel (ET700/75m, Chroma) was recorded for 2 min. Data was analyzed by a home-made MATLAB script.

### Box-plot

The function of “boxplot” in MATLAB software (R2015a, 64-bit, February 12, 2015) was used to plot the boxplots in Supplementary Fig. 2 and 4. For each boxplot, the red bar represents median. The bottom edge of the box represents 25th percentiles, and the top is 75th percentiles. Most extreme data points are covered by the whiskers except outliers. The ‘+’ symbol is used to represent the outliers.

### Unpaired t test

Statistical significance was evaluated based on Student’s t-tests (Prism 9 for macOS, Version 9.1.0 (216), March 15, 2021, GraphPad Software, Inc.). Test was chosen as unpaired t test. P value style: GP: 0.1234 (ns), 0.0332 (*), 0.0021 (**), 0.0002 (***), < 0.0001 (****).

### Dimensionless generic model and numerical simulation

The dynamical model (Eq. 1) can be rescaled to the following general system and expressed in terms of rescaled *k*_*d*_ = 1 and *D*_*v*_ = 1 (see Supplementary texts 1.4 for details)

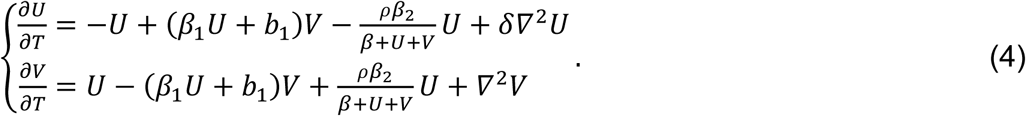

The interpretation of the parameters in this nondimensionalized model (Eq. 4), their default values, and their definition in terms of the original experimental measurements in our study are given in Table 1 and Supplementary Table 1. As Eq. 4 might prove general and unspecific, we conducted numerical simulations based on the dimensionless system (Eq. 4) to unify MCRD systems.

