## Supplemental files for "Biomolecular condensates bridge experiment and theory of mass-conserving reaction-diffusion systems in phase separation"

### Supplementary Information

Table of contents

**1. Supplementary Methods**

**1.1 Description of the mass conservation dynamical model**

**1.2 Numerical simulation**

**1.3 Spatial scale and structure factors**

**1.4 Estimating the dispersion relation of phase-separation emergence**

**1.5 Obtaining dispersion diagrams from experiment data**

**1.6 Analysis of eigenvalue function**

**2. Protein and dsDNA sequences**

**3. Supplementary Figures**

**4. Supplementary Tables**

**5. Supplementary Movie Legends**

#### 1. Supplementary Methods

Our previous investigation (*47*) has demonstrated that a truncated version of the human transcription factor p53, referred to as p53$^{4M}$ ΔTAD, in conjunction with dsDNA, can give rise to DPICs (Fig. 1A). This mutant lacks the amino-terminal transactivation domain (TAD, 1-62) and contains mutations in four residues (M133L/V203A/N239Y/N268D) within the dsDNA-binding domain (DBD) (Supplementary Fig. 1A & **Methods**). Following the protocol outlined in previous study, we purified the p53 variant "p53$^{4M}$ ΔTAD" *in vitro* (Supplementary Fig. 1B) and conducted electrophoretic mobility shift assays (EMSAs) to confirm the dsDNA-binding affinity and sequence selectivity (Supplementary Fig. 1C), validating its activity.

Upon mixing 20 μM p53$^{4M}$ ΔTAD labeled with ATTO565 and 0.6 μM 400-bp random dsDNA labeled with FAM for 0.5 hours, droplet-like DPICs formed (Fig. 1A). To scrutinize the intricate details of DPICs, we conducted *in vitro* droplet assays, systematically combining concentrations of 0, 0.15, 0.3, 0.6, and 0.9 μM 400-bp random dsDNA labeled with DAPI with 0, 2.5, 5, 10, 20, and 40 μM dark p53$^{4M}$ ΔTAD, thereby constructing a phase diagram (Supplementary Fig. 1D). It is pertinent to mention that the working buffer employed in our experiments maintained the physiological conditions (8 mM Tris-HCl (pH 7.5) and 120 mM NaCl) and contained no crowding agents. Furthermore, all referenced p53 concentrations pertain to the monomeric form.

##### 1.1 Description of the mass conservation dynamical model

We then investigated whether the DPIC system exhibits local concentration-dependent feedback mechanisms, which are crucial for establishing the mass-conserving dynamical model. Throughout the formation process of DPICs, the total amount of free protein and dsDNA-bound protein remains conserved. This conservation also applies to the dsDNA. Previous fluorescence recovery after photobleaching (FRAP) experiments have demonstrated that the motility of long dsDNA is significantly lower than that of proteins inside the DPIC (Supplementary Fig. 2A), and other methods demonstrated that the DPIC formation comes from the capacity of p53 to bridge different dsDNA duplexes (*47*). Additionally, fluorescence correlation spectroscopy (FCS) has shown that the diffusion coefficient of dsDNA is lower than that of proteins outside the DPIC (Supplementary Fig. 2B). Therefore, we mainly focused on the change of protein concentration inside and outside of DPIC.

In our *in vitro* DPIC experiments (Supplementary Fig. 1), we observed two distinct phases: inside DPIC and outside DPIC, as depicted in our microscopy images (Figs. 1A(i) and 2B(iii)-(iv)). Additionally, the onset of these two phases depends on the initial protein concentration within the system. Time-lapse videography revealed that the droplet-like patterns exhibit coarsening dynamics. Utilizing the previous work, we verified the DPIC is a network-like structure by protein bridging different dsDNA duplexes (*47*), so we identify two states of protein p53$^{4M}$ ΔTAD, dsDNA-bound state and the free state. The emergence of the droplet DPICs from spatial homogenous into two distinct phases due to redistribution processes of these two states of proteins. Consequently, we developed a MCRD model that incorporates the concentrations of dsDNA-bound protein $u(x, y, t)$ and free protein $v(x, y, t)$, enabling us to explore the underlying mechanisms of phase-separation pattern formation, with $(x, y)$ and $t$ representing space and time, respectively. The dynamics of droplet formation can be delineated by three fundamental processes, based on the principle of mass conservation: Spontaneous dissociation (*49*), recruitment (Supplementary Fig. 3), and passive escape from DPIC (Supplementary Fig. 5). The buffer solution serves as hosts for the unbounded protein.

Both experimental findings and molecular dynamics simulations demonstrate that dsDNA-bound proteins can undergo spontaneous dissociation, transitioning into free proteins within the DPIC (*47*). We have quantitatively analyzed the correlation between the initial normalized intensity of the dsDNA-bound protein concentration and the sequestration of free proteins (see Fig. 1B(i)-(ii) and Supplementary Fig. 3). The rate of recruitment is detailed in the results section (see Fig. 1B(ii)), and the corresponding mathematical model is as follows (Supplementary Table 2 for model selection):

$$f(u) = k_1 u + b,$$

where $u$ represents the concentration of dsDNA-bound protein, while $b$ denotes the constant rate of spontaneous aggregation in the absence of a dsDNA-bound cluster. Consequently, the recruitment rate is directly proportional to both the concentration of free protein and the concentration of the cluster, as detailed in Supplementary Fig. 3.

Finally, our experiments using 10 μM p53$^{4M}$ ΔTAD and 0.3 μM 400-bp random dsDNA have identified an escape process. In this process, dsDNA-bound proteins transition into outside the DPIC from the inside DPIC (Fig. 1C(i)-(ii) and Supplementary Fig. 5). The reduction in protein concentration is influenced by both dsDNA characteristics—specifically length and sequence—and the combined concentration of dsDNA-bound and free proteins $(u + v)$. For simplicity, we introduce the parameter $\rho$ to represent the stability of arrest protein, accounting for variations in length or sequence. Our theoretical model and experimental data, suggesting a nonlinear decay described by the following formula (Supplementary Table 2 for model selection):

$$g(u, v) = \frac{\beta}{u+v+\beta},$$

where $\beta$ is the half-saturation constant to regulate the effect of dsDNA-bound protein on escape rate.

The mathematical model integrates the three processes previously mentioned, providing a comprehensive representation of the biomolecular condensates. The concentration of dsDNA-bound protein can be effectively estimated using the following reaction-diffusion equations:

$$\begin{cases} \frac{\partial u}{\partial t} = \underbrace{-k_d u + f(u)v - \rho k_2 g(u,v)u}_{\mathcal{F}(u,v)} + D_u \nabla^2 u, \\ \frac{\partial v}{\partial t} = k_d u - f(u)v + \rho k_2 g(u,v)u + D_v \nabla^2 v. \end{cases} \quad \text{(S1)}$$

In model (Eq. S1), the reaction function $\mathcal{F}(u,v)$ describes the dynamic transition of proteins from a free to a dsDNA-bound state. The first term illustrates the spontaneous dissociation of dsDNA-bound proteins into free proteins. The second and third terms respectively quantify the recruitment and escape processes, which are both concentration-dependent feedback. Specifically, they are proportional to the concentrations of free protein ($v$) and dsDNA-bound protein ($u$). The parameter $\rho$ quantifies the stability of dsDNA-bound arrested proteins during the escape processes and is expected to be positive, reflecting the influence of energy barriers and surrounding temperature ($k_B T$). The diffusion coefficients are empirical constants, with the constraint that $D_u < D_v$ reflecting the relative Brownian movement of the components within the buffer (Supplementary Fig. 2). The detailed definition, dimensions, and values of the model parameters are provided in Supplementary Table 1. The solutions of this model (Eq. S1) are discussed in the numerical simulation section as follows.

#### 1.2 Numerical simulation

Model simulations were executed in two-dimensional space by discretizing $t, x,$ and $y$ in jupyterlab with PyOpenCL using finite difference methods with periodic boundary conditions and non-dimensional units. The change of $u, v$ and the diffusive fluxes were calculated by forward differences and second-order central differences, respectively. Different grid sizes and physical lengths were applied for numerical simulations to confirm robustness of our results. We adopted periodic boundary condition for all the simulations except for the phase diagram with Neumann boundary in Fig. 2A. Computational tasks were offloaded onto Quadro K5200 to expedite the processing speed. Initial simulation conditions consisted of a homogeneous distribution of $u$ and $v$ with a slight random perturbation. All simulations were derived with a timestep of $\Delta t$ = 0.02 s and spatial discretization of $\Delta x$ = 0.2. The detailed parameters are listed in Table 1, and the code is available on the github, https://github.com/liuqx315/DPICs-MCRD-Phase-separation.

#### 1.3 Spatial scale and structure factors

**Spatial scale.** Quantitative analysis of wavelength was conducted to determine the spatial scale of the patterns observed both in the experimental and simulation images. For simulation images, we performed a 2D Fast Fourier Transform to obtain the power spectral density (PSD) in a square window and compute the wavelength by identifying the maximum PSD. For experimental images exhibiting noticeable noise, Square, moving windows were applied for determining local wavelengths, and the results were then averaged across all identified windows. This effective technique for calculating wavelength in noisy images is described in the previous references (*20, 69*).

**Structure factors.** Structural factor analysis, a commonly utilized approach for investigating spatiotemporal self-similar dynamic properties, typically holds a robust scaling law. The dynamic structure factor $\boldsymbol{S}(\boldsymbol{q},\boldsymbol{t})$ is derived by performing the 2D Fourier transform of the binarized experimental and simulation images. It is defined as $\boldsymbol{S}(\boldsymbol{q},\boldsymbol{t}) = \langle |\hat{\boldsymbol{\phi}}(\boldsymbol{q},\boldsymbol{t})|^{\mathbf{2}} \rangle$ (*50*), where $\hat{\phi}(\boldsymbol{q},t)$ represents the Fourier transform and $\boldsymbol{q}$ denotes wave vector. The circularly averaged structure factor $S(q,t)$ defined as $S(q,t) = \langle |\hat{\phi}(\boldsymbol{q},t)|^2 \rangle_q$, with $q$ being the modulus of $\boldsymbol{q}$, is generated by computing the Modulus squared of the aforementioned Fourier transform. Then we extract the wave-vector magnitude $q_{max}$ by calculating $q_{max}(t) = \frac{\int_0^\infty qS(q,t)dq}{\int_0^\infty S(q,t)dq}$.

##### 1.4 Estimating the dispersion relation of phase-separation emergence

**Derivation of the theoretical dispersion relation.** The theoretical line of dispersion relation is derived by qualitatively plot of the maximal eigenvalue of the wave numbers. The primary approach involved linearizing the full system to obtain an eigenvalue problem. To achieve this, we first apply the following rescaled relations to obtain the general dimensionless model presented below.

**(1) Rescaled parameters of the model** (**S1**). Let $U = u, V = v, \delta = \frac{D_u}{D_v}, \beta_1 = \frac{k_1}{k_d}, \beta_2 = \frac{k_2\beta}{k_d}, T = k_d t, b_1 = \frac{b}{k_d}, \beta = \beta, \rho = \rho, X = k_d^{\frac{1}{2}} D_v^{\frac{-1}{2}} x$, then we obtain the model as follows,

$$\begin{cases} \frac{\partial U}{\partial T} = -U + (\beta_1 U + b_1)V - \frac{\rho\beta_2}{\beta + U + V} U + \delta \nabla^2 U \\ \frac{\partial V}{\partial T} = U - (\beta_1 U + b_1)V + \frac{\rho\beta_2}{\beta + U + V} U + \nabla^2 V \end{cases}, \quad \text{(S2)}$$

where $\delta < 1.0$ for biological meaning. Upon the dimensionless model, neglect the abuse of notation and we get the expressions,

$$U = u, V = v, f(u) = (\beta_1 u + b_1), g(u,v) = \frac{\beta_2}{\beta + u + v},$$

and

$$\mathcal{F}(u,v) = -u + f(u)v - \rho g(u,v)u.$$

**(2) Linearization and eigenvalues**

The nonspatial system (S2) has one stable state when we neglect spatial coupling. We denote the fixed point as $(\bar{u},\bar{v})$.

Substitute $u = \bar{u} + \hat{u}e^{\sigma t + i\boldsymbol{q}\vec{r}}$, $v = \bar{v} + \hat{v}e^{\sigma t + i\boldsymbol{q}\vec{r}}$ into system (S2) and implement a Taylor expansion around $(\bar{u},\bar{v})$, we obtain the eigenvalue equation,

$$\sigma\binom{\hat{u}}{\hat{v}} = \begin{pmatrix} \mathcal{F}_u - \delta q^2 & \mathcal{F}_v \\ -\mathcal{F}_u & -\mathcal{F}_v - q^2 \end{pmatrix}\binom{\hat{u}}{\hat{v}} \stackrel{\text{def}}{=} A\binom{\hat{u}}{\hat{v}}. \quad \text{(S3)}$$

The corresponding characteristic polynomial is given as

$$\sigma^2 + \sigma\big(\mathcal{F}_v - \mathcal{F}_u + q^2(\delta+1)\big) + q^2(\delta\mathcal{F}_v - \mathcal{F}_u) + \delta q^4 = 0. \quad \text{(S4)}$$

Subsequently, we obtain the eigenvalues of system (S2):

$$\lambda_{1,2} = \frac{1}{2}tr(A) \pm \frac{1}{2}\sqrt{\mathrm{tr(A)}^2 - 4\mathrm{det(A)}}. \quad \text{(S5)}$$

where

$$\mathrm{tr}(A) = \mathcal{F}_u - \mathcal{F}_v - q^2(\delta+1) = \beta_1 v - \beta_1 u - b_1 - q^2(\delta+1) - \frac{\beta_2\rho}{\beta+u+v} - 1, \quad \text{(S6)}$$

$$\begin{aligned}\det(A) &= q^2(\delta\mathcal{F}_v - \mathcal{F}_u + \delta q^2) \\ &= q^2\left(\delta\left(b_1 + \beta_1 u + \frac{\beta_2\rho u}{(\beta+u+v)^2}\right) - \beta_1 v + \delta q^2 + \frac{\beta_2\rho}{\beta+u+v} - \frac{\beta_2\rho u}{(\beta+u+v)^2} + 1\right). \quad \text{(S7)}\end{aligned}$$

Since $Re(\sigma_1) > Re(\sigma_2)$, the theoretical line is therefore obtained as a function of real part of the maximal eigenvalue $Re(\sigma_1)$ associated with perturbation wavenumber $q$, with $\bar{u} = 16.6, \bar{v} = 3.9, \beta_1 = 3.75, \beta_2 = 100, b_1 = 25.0, \beta = 2.0, \rho = 4.0, \delta = 0.003$ (red solid line in Fig. 4b), $\bar{u} = 35.1, \bar{v} = 2.7, \beta_1 = 3.75, \beta_2 = 100, b_1 = 25.0, \beta = 2.0, \rho = 16.0, \delta = 0.0005$ (brown solid line in Fig. 4b), respectively.

#### 1.5 Obtaining dispersion relation from experimental data

In previous self-organized patterns, only theoretical dispersion relations have been calculated within textbooks (*70*). To date, there is still no methodology available for obtaining experimental dispersion relations. Based on theoretical analysis, we observe fundamental differences between the classical Turing theory and phase-separation principles concerning dispersion relations (*9, 70*). Here, we describe how we measure the growth rate of phase-separation patterns as a function of their wave numbers $q$ from successive frames throughout the duration of our DPIC experiment.

The methodology for obtaining dispersion relation from experimental data is based on the assumptions of that the initial growth rate of each mode (wavelength) are independent of each other. Obviously, the negative and positive growth rates are particularly important. Following our linear stability analysis (Eq. S2), we know that the amplitude $r(q,t)$ is the exponential growth with time described by formular

$$r(q,t) \propto e^{\sigma(q)t}. \qquad \text{(S8)}$$

Here, $\sigma(q)$ denotes the corresponding growth rate with respect to wave number $q$, during the linear growth phase. In this expression, $q = \frac{2\pi}{\lambda}$ is the wave number and $\sigma(q)$ the growth rate with unites of frequency. Positive and negative growth rates correspond to unstable and stable modes, respectively.

The process encompasses 2 main steps as follows:

**2D Spectral Analysis of Experiment Images.** We perform 2D spectral analysis of experiment images from time period 300s ~ 600s with each image taken at intervals of 30s (i.e.11 images) for experiment group 10 μM $p53^{4M}$ ΔTAD and 0.3 μM 400-bp random dsDNA. This time period is considered within linear regime. And for experiment group 20 μM $p53^{4M}$ ΔTAD and 0.6 μM 400-bp random dsDNA the linear regime is considered to be 120s ~300s (i.e. 7 images). For each image, an algorithm based on Fourier analysis is applied to extract radial amplitudes of different frequency components, or equivalently, each isolated wavenumber $q$. This approach indirectly computes the Fourier transform by traversing each pixel of the image and calculating the weighted sum of cosine and sine functions at various frequencies. The amplitude of each frequency component is determined by taking the square root of the sum of squares of the cosine and sine components. Consequently, a sequence of radial amplitudes $r(q)$ corresponding to different wavenumbers is obtained.

**Exponential Growth Analysis.** For each $q$, a time sequence of radial amplitude evolution is obtained. Leveraging on the exponential growth assumption $r(q,t) \propto exp(\sigma(q)t)$ within linear regime, we take logarithmic transformation of $r(q,t)$ and conduct linear regression analysis using *fitlm* function in Matlab R2023a to determine the corresponding exponential coefficient $\sigma(q)$, and the intercept $b(q)$. Therefore, we have computed the growth rate $\sigma(q)$ with respect to different $q$. In other words, we obtain the dispersion diagram from experiment data.

Note that in Fig. 4, the wavenumber is normalized by the length of image with 184 $\mu$m by 184 $\mu$m. And the experimentally derived growth rate $\sigma(q)$ scales as $\frac{\sigma(q)}{C}$, where $C$ is a scaling constant chosen as $C = \frac{max(Re(\lambda_1))}{max(\sigma(q))}$ in our case.

#### 1.6 Analysis of eigenvalue function

$$\frac{\partial U}{\partial T} = -U + (\beta_1 U + b_1)\cdot V - \rho\cdot\left(\frac{\beta_2}{\beta+U+V}\right)\cdot U + \delta\nabla^2 U \quad \text{(S9a)}$$

$$\frac{\partial V}{\partial T} = U - (\beta_1 U + b_1)\cdot V + \rho\cdot\left(\frac{\beta_2}{\beta+U+V}\right)\cdot U + \nabla^2 V \quad \text{(S9b)}$$

Where $\delta \ll 1.0$,for biological meaning. Upon the dimensionless model, neglect the abuse of notation and we get the expressions,

$$U = u, V = v, f(u) = \beta_1 u + b_1, g(u,v) = \frac{\rho\beta_2}{\beta+u+v},$$

$$\mathcal{F}(u,v) = -u + f(u)v - g(u,v)u,$$

$$\mathcal{F}_u = \beta_1 v - \frac{\beta_2\rho}{\beta+u+v} + \frac{\beta_2\rho u}{(\beta+u+v)^2} - 1,$$

$$\mathcal{F}_v = b_1 + \beta_1 u + \frac{\beta_2\rho u}{(\beta+u+v)^2}.$$

The corresponding characteristic polynomial around steady state$(\bar{u},\bar{v})$is given as,

$$\sigma^2 + \sigma(\mathcal{F}_v - \mathcal{F}_u + q^2(\delta+1)) + q^2(\delta\mathcal{F}_v - \mathcal{F}_u) + \delta q^4 = 0. \quad \text{(S10)}$$

Subsequently, we obtain the eigenvalues of system (S9):

$$\sigma_{1,2} = \frac{1}{2}\left(tr(A) \pm \sqrt{tr(A)^2 - 4det(A)}\right),$$

Where

$$tr(A) = \mathcal{F}_u - \mathcal{F}_v - q^2(\delta+1) = \beta_1 v - \beta_1 u - b_1 - q^2(\delta+1) - \frac{\beta_2\rho}{\beta+u+v} - 1, \quad \text{(S11)}$$

$$det(A) = q^2(\delta\mathcal{F}_v - \mathcal{F}_u + \delta q^2)$$

$$= q^2(\delta(b_1 + \beta_1 u + \frac{\beta_2\rho u}{(\beta+u+v)^2}) - \beta_1 v + \delta q^2 + \frac{\beta_2\rho}{\beta+u+v} - \frac{\beta_2\rho u}{(\beta+u+v)^2} + 1. \quad \text{(S12)}$$

To compute the expression for $\sigma_1$, one should note that we substitute steady state value $\bar{u},\bar{v}$ into matrix $A$.

$$\sigma_1 = \frac{1}{2}(\beta_1\overline{v} - \beta_1\overline{u} - b_1 - \eta_3 - 1) - (\frac{1}{2}\frac{\beta_2\rho}{\beta+\overline{u}+\overline{v}}) + \frac{1}{2}\sqrt{(b_1+\beta_1\overline{u}-\beta_1\overline{v}+\eta_3+\eta_2+1)^2 - 4q^2(\delta(b_1+\beta_1\overline{u}+\eta_1) - \beta_1\overline{v} + \delta q^2 + \eta_2 - \eta_1 + 1)}, \tag{S13}$$

where

$$\eta_1 = \frac{\beta_2\rho\overline{u}}{(\beta+\overline{u}+\overline{v})^2}, \tag{S14}$$

$$\eta_2 = \frac{\beta_2\rho}{(\beta+\overline{u}+\overline{v})}, \tag{S15}$$

$$\eta_3 = q^2(\delta+1). \tag{S16}$$

Denote $C_3 = \frac{1}{2}(b_1+\beta_1\overline{u}-\beta_1\overline{v}+\eta_3+\eta_2+1)^2$, $C_3$ is a variable constant that relies on $\beta_1, \overline{u}, \overline{v}, b_1, q, \delta$ but independent of $\rho$.

Now consider the square root term.

Let $B = (b_1+\beta_1\overline{u}-\beta_1\overline{v}+\eta_3+\eta_2+1)^2$, and denote $C_1(b_1,\beta_1,\overline{u},\overline{v},b_1,q,\delta) = 1 + b_1 + \beta_1\overline{u} - \beta_1\overline{v} + \eta_3$. Note that $C_1$ is also a variable constant independent of $\rho$. Then $B = (C_1+\eta_2)^2 = C_1^2 + 2C_1\eta_2 + \eta_2^2$. Trying to find out the relation between $\sigma_1$ and $\rho$, we extract the terms related to $\rho$. Therefore, $B$ contributed two entries related to $\rho$, that is,

$$2C_1\frac{\beta_2\rho}{(\beta+\overline{u}+\overline{v})^2} + \frac{\beta_2^2\rho^2}{(\beta+\overline{u}+\overline{v})^4}.$$

Let $C = -4q^2(\delta(b_1+\beta_1\overline{u}+\eta_1) - \beta_1\overline{v} + \delta q^2 + \eta_2 - \eta_1 + 1)$.
Denote $C_2 = -4q^2(\delta b_1 + \delta\beta_1\overline{u} - \beta_1\overline{v} + \delta q^2 + \eta_2 - \eta_1 + 1) = C_2(\delta, b_1, \beta_1, \overline{u}, \overline{v}, b_1, q)$. The $C$ can be written as

$$C = C_2 - 4q^2(\delta\eta_1 + \eta_2 - \eta_1).$$

Consider the term $-4q^2(\delta\eta_1+\eta_2-\eta_1)$,

$$-4q^2(\delta\eta_1+\eta_2-\eta_1) = q^2\eta_1(1-\delta) - 4q^2\eta_2 \tag{S17}$$

$$= 4q^2\left[\left((1-\delta)\frac{\beta_2\overline{u}}{(\beta+\overline{u}+\overline{v})^2}\right)\cdot\rho - \frac{\beta_2\rho}{(\beta+\overline{u}+\overline{v})}\right] \tag{S18}$$

$$= \frac{4q^2\beta_2}{(\beta+\overline{u}+\overline{v})}\left[\frac{(1-\delta)}{(\beta+\overline{u}+\overline{v})-1}\right]\cdot\rho. \tag{S19}$$

Hence,

$$\sigma_1 = C_3 - \left(\frac{1}{2}\frac{\beta_2\rho}{\beta + \bar{u} + \bar{v}}\right) + \frac{1}{2}\sqrt{C_1^2 - C_2 + \mathcal{G}(\rho)}.$$

Where

$$\mathcal{G}(\rho) = \frac{\beta_2^2\rho^2}{(\beta + \bar{u} + \bar{v})^2} + \frac{\beta_2\rho}{(\beta + \bar{u} + \bar{v})} \cdot \left(\frac{4q^2(1-\delta)\bar{u} + 2C_1}{\beta + \bar{u} + \bar{v}} - 4q^2\right) \tag{S20}$$

$$= \frac{\beta_2\rho}{(\beta + \bar{u} + \bar{v})}\left(\frac{\beta_2\rho}{(\beta + \bar{u} + \bar{v})^3} + \frac{-4q^2\delta\bar{u} + 2C_1 - 4q^2\beta - 4q^2\bar{v}}{\beta + \bar{u} + \bar{v}}\right) \tag{S21}$$

$$= \frac{\beta_2\rho}{(\beta + \bar{u} + \bar{v})} \cdot \left[\frac{\beta_2\rho}{(\beta + \bar{u} + \bar{v})^3} + \frac{-4q^2(\delta\bar{u} + \beta + \bar{v}) + 2C_1}{\beta + \bar{u} + \bar{v}}\right]. \tag{S22}$$

In our proposed model, the effect of dsDNA on this system relies on parameter $\rho$. To see how $\rho$ contributes to $\sigma_1$, we then assume $\beta$, $\beta_1$, $\beta_2$, $b_1$ are constants with default value mentioned in Table 1 and neglect sufficiently small terms. In reality, $\delta \approx o(10^{-3})$, and perturbation $q$ is small. Therefore, we have the following simplification.

$$C_3 = \frac{15}{8}\big((\bar{v} - \bar{u}) - q^2(\delta + 1)\big) - \frac{15}{4} \approx \frac{15}{8}\big((\bar{v} - \bar{u}) - q^2\big) - \frac{15}{4},$$

$$C_1 = 26 + 3.75(\bar{u} - \bar{v}) + q^2(\delta + 1) \approx 26 + 3.75(\bar{u} - \bar{v}) + q^2,$$

$$C_2 = -4q^2(25\delta + \delta 3.75(\bar{u} - \bar{v}) + \delta q^2 + 1) \approx 15q^2\bar{v} - 4q^2,$$

$$\mathcal{G}(\rho) \approx \frac{\beta_2\rho}{\beta + \bar{u} + \bar{v}}\left[\frac{\beta_2\rho}{(\beta + \bar{u} + \bar{v})^3} + \frac{-4q^2(\beta + \bar{v} + 2C_1)}{\beta + \bar{u} + \bar{v}}\right],$$

$$\frac{1}{2}\sqrt{C_1^2 - C_2 + \mathcal{G}(\rho)} \approx q^4 + 4q^2 - 15q^2\bar{v} + 728 + 202(\bar{u} - \bar{v}) + \frac{15^2}{4}(\bar{u} - \bar{v})^2 \tag{S23}$$

$$= \frac{\beta_2\rho}{(\beta+\bar{u}+\bar{v})}\left(\frac{\beta_2\rho}{(\beta+\bar{u}+\bar{v})^3} + \frac{-4q^2\delta\bar{u}+2C_1-4q^2\beta-4q^2\bar{v}}{\beta+\bar{u}+\bar{v}}\right) \tag{S24}$$

$$= \frac{\beta_2\rho}{(\beta+\bar{u}+\bar{v})} \cdot \left[\frac{\beta_2\rho}{(\beta+\bar{u}+\bar{v})^3} + \frac{-4q^2(\delta\bar{u}+\beta+\bar{v})+2C_1}{\beta+\bar{u}+\bar{v}}\right], \tag{S25}$$

$$C_3 - \frac{1}{2}\frac{\beta_2\rho}{\beta + \bar{u} + \bar{v}} \approx -\left(\frac{15}{8}(\bar{u} - \bar{v}) + q^2 + \frac{1}{2}\frac{\beta_2\rho}{\beta + \bar{u} + \bar{v}}\right)\Bigg) := -K(\rho).$$

Hence, Eq. (S23) can be rewritten as

$$\frac{1}{2}\sqrt{C_1^2 - C_2 + \mathcal{G}(\rho)} \approx \mathrm{K}(\rho) \cdot \sqrt{1 + \left(\frac{\beta_2\rho}{(\beta+\overline{u}+\overline{v})^2}\right)^2 + D_1 + D_2}, \quad \text{(S26)}$$

where

$$D_1 = \frac{1}{(\beta + \overline{u} + \overline{v})^2}\left(-7q^2 + 4q^2\overline{v} + 52 + 7.5(\overline{u} - \overline{v})\right),$$

$$D_2 = -15q^2\overline{v} - 3q^4 + 4q^2 + 728 + 202(\overline{u} - \overline{v}).$$

Therefore

$$\sigma_1 = \mathrm{K}(\rho)\left[\sqrt{1 + \left(\frac{\beta_2\rho}{(\beta+\overline{u}+\overline{v})^2}\right)^2 + \frac{\beta_2\rho}{(\beta+\overline{u}+\overline{v})^2} - \frac{\beta_2\rho}{\beta+\overline{u}+\overline{v}}\frac{15q^2}{2}(\overline{u} - \overline{v}) + D_1 + D_2} - 1\right]. \quad \text{(S27)}$$

It is straightforward to verify that $\rho$ exists only on the denominator of the expression of $\sigma_1$ in Eq. (S27). There are two observations can be deduced when increasing $\rho$ only: one is that $\sigma_1$ will decrease for a fixed wavenumber $q$, and the other is that the dispersion relation will cross the x-axis later.

#### 2. Supplementary Methods

##### 2.1 Protein sequence

| Protein | Sequence |
|---|---|
| 6× His-p53$^{4M}$ ΔTAD<br>(p53$^{4M}$ ΔTAD) | MGSSHHHHHHAMAAPRMPEAAPRVAPAPAAPTPAAPAPAPSWPL<br>SSSVPSQKTYQGSYGFRLGFLHSGTAKSVTCTYSPALNKLFCQLA<br>KTCPVQLWVDSTPPPGTRVRAMAIYKQSQHMTEVVRRCPHHERC<br>SDSDGLAPPQHLIRVEGNLRAEYLDDRNTFRHSVVVPYEPPEVGS<br>DCTTIHYNYMCYSSCMGGMNRRPILTIITLEDSSGNLLGRDSFEVR<br>VCACAGRDRRTEEENLRKKGEPHHELPPGSTKRALPNNTSSSPQ<br>PKKKPLDGEYFTLQIRGRERFEMFRELNEALELKDAQAGKEPGGS<br>RAHSSHLKSKKGQSTSRHKKLMFKTEGPDSD* |

##### 2.2 dsDNA sequences (the top strand was shown)

| Application of dsDNA | Sequence (5' to 3'), p21 sequence shown in red |
|---|---|
| 30-bp random dsDNA for EMSA | AAC CTG TCG TGC CAG CTG CAT TAA TGA ATC |
| 30-bp dsDNA with 1× p21 binding motif for EMSA | TCT AT GAA CAT GTC CCA ACA TGT TG TTC CC |
| 400-bp random dsDNA for *in vitro* droplet assay | CTT TTG CTT GAT CTC AGT TTC AGT ATT AAT ATC CAT TTT<br>TTA TAA GCG TCG ACG GCT TCA CGA AAC ATC TTT TCA<br>TCG CCA ATA AAA GTG GCG ATA GTG AAT TTA GTC TGG<br>ATA GCC ATA AGT GTT TGA TCC ATT CTT TGG GAC TCC<br>TGG CTG ATT AAG TAT GTC GAT AAG GCG TTT CCA TCC<br>GTC ACG TAA TTT ACG GGT GAT TCG TTC AAG TAA AGA TTC<br>GGA AGG GCA GCC AGC AAC AGG CCA CCC TGC AAT GGC |

| | |
|---|---|
| | ATA TTG CAT GGT GTG CTC CTT ATT TAT ACA TAA CGA AAA ACG CCT CGA GTG AAG CGT TAT TGG TAT GCG GTA AAA CCG CAC TCA GGC GGC CTT GAT AGT CAT ATC ATC TGA ATC AAA TAT TCC TGA TGT ATC GAT ATC GGT A |

#### 3. Supplementary Figures

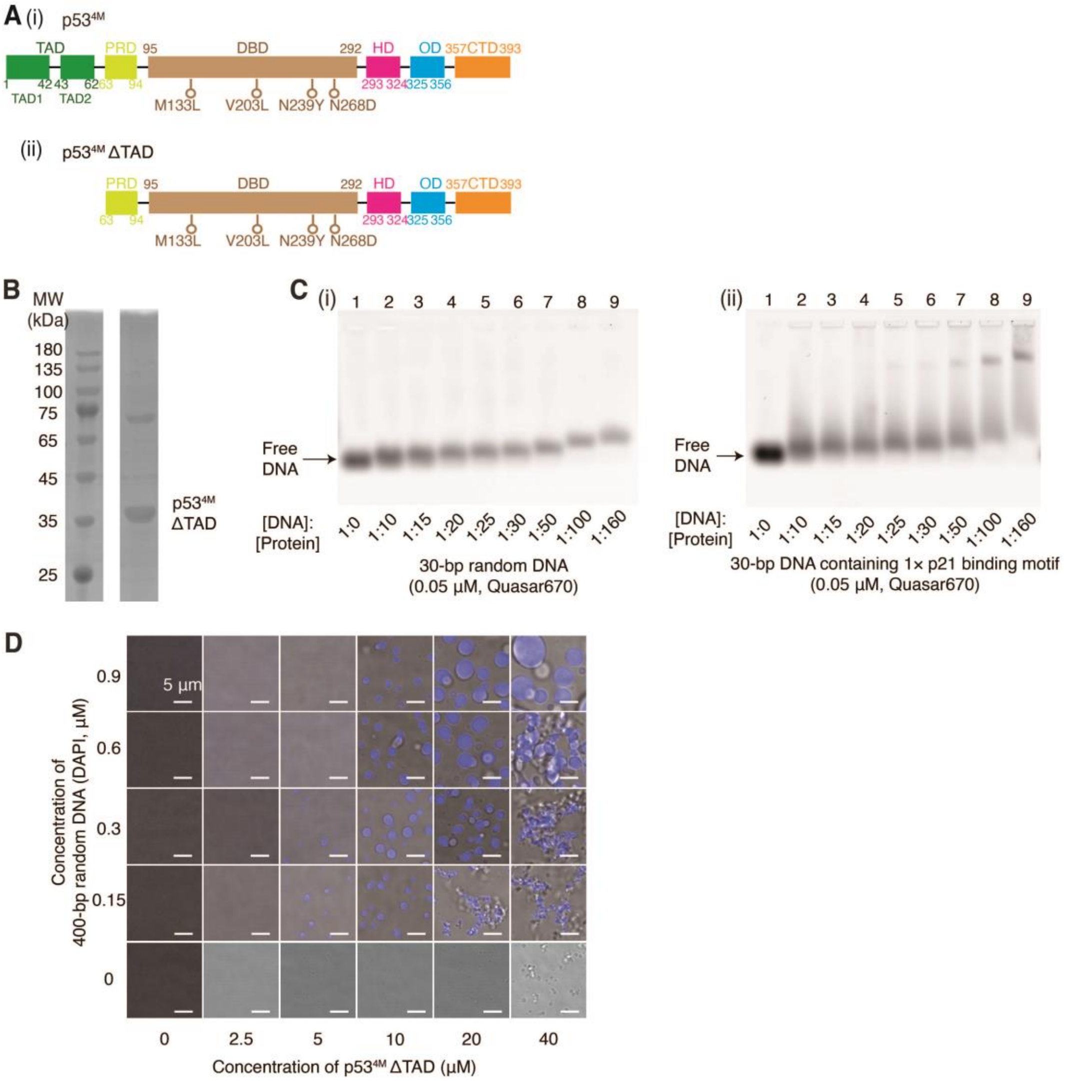


**Supplementary Fig. 1. *In vitro* purified p53$^{4M}$ ΔTAD is active.** (**A**) Schematic diagram of p53$^{4M}$ and p53$^{4M}$ ΔTAD. (**B**) SDS-PAGE analysis of p53$^{4M}$ ΔTAD. (**C**) EMSAs (1% agarose gel) showing p53$^{4M}$ ΔTAD interaction with 30-bp dsDNA. (i) 30-bp random dsDNA; (ii) 30-bp dsDNA containing 1× p21 binding motif. dsDNA substrates were labeled by Quasar670, and imaged by an Amersham Typhoon RGB system (with a 635 nm laser and Cy5 670BP30 filter). (**D**) *In vitro* droplet assays mixing 0, 0.15, 0.3, 0.6, and 0.9 μM 400-bp random dsDNA with 0, 2.5, 5, 10, 20, and 40 μM dark p53$^{4M}$ ΔTAD, where dsDNA was labeled with DAPI. All reactions were incubated at room temperature for 30 minutes. The working buffer was 8 mM Tris-HCl (pH 7.5), 120 mM NaCl, 4% Glycerol, and 16 mM DTT, without any crowding agents.

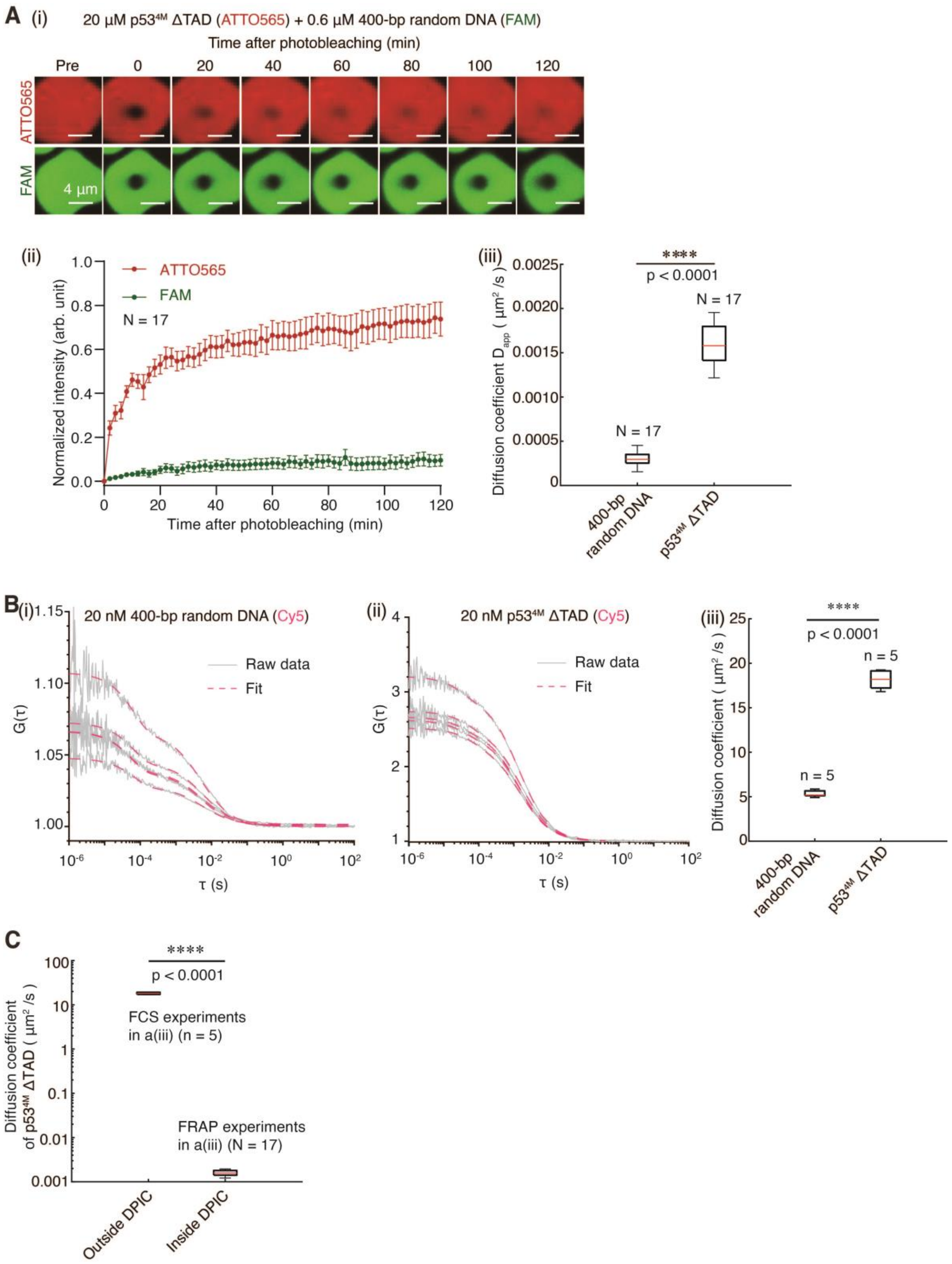


**Supplementary Fig. 2. The measurement of the diffusion coefficient of p53$^{4M}$ ΔTAD inside and outside the DPICs.** (**A**) The FRAP experiments were conducted to measure the diffusion coefficients of dsDNA and protein inside the DPICs. (i) DPICs formed by 0.6

μM 400-bp random dsDNA labeled with FAM and 20 μM p53$^{4M}$ ΔTAD labeled with ATTO565, with a 2-hour incubation and 2-hour recovery process recorded. (ii) FRAP curves with red indicating p53$^{4M}$ ΔTAD (ATTO565) and green indicating dsDNA (FAM). Seventeen independent DPICs were plotted (N = 17). Error bars represent mean ± s.d. (i) and (ii) were shown in the previous work (*47*). (iii) Apparent diffusion coefficient ($D_{app}$) of 400-bp random dsDNA and p53$^{4M}$ ΔTAD inside DPIC. Unpaired t test was performed. (**B**) The FCS experiments were conducted to measure the diffusion coefficients of dsDNA and protein outside the DPICs. (i) FCS curves for 20 nM 400-bp random dsDNA. (ii) FCS curves for 20 nM p53$^{4M}$ ΔTAD. The working buffer for FCS assays was 8 mM Tris-HCl (pH 7.5), 120 mM NaCl, and 16 mM DTT. Independent FCS assays were performed 5 times for this condition (n = 5). (iii) Diffusion coefficients of free 400-bp random dsDNA and p53$^{4M}$ ΔTAD. Unpaired t test was performed. (**C**) Comparison of the diffusion coefficient of p53$^{4M}$ ΔTAD outside the DPIC and that inside DPIC. Unpaired t test was performed. For the boxplot, the red bar represents median. The bottom edge of the box represents 25$^{th}$ percentiles, and the top is 75$^{th}$ percentiles. Most extreme data points are covered by the whiskers except outliers. The '+' symbol is used to represent the outliers. Statistical significance was analyzed using unpaired t test for two groups. P value: two-tailed; p value style: GP: 0.1234 (ns), 0.0332 (*), 0.0021 (**), 0.0002 (***), <0.0001 (****). Confidence level: 95%.

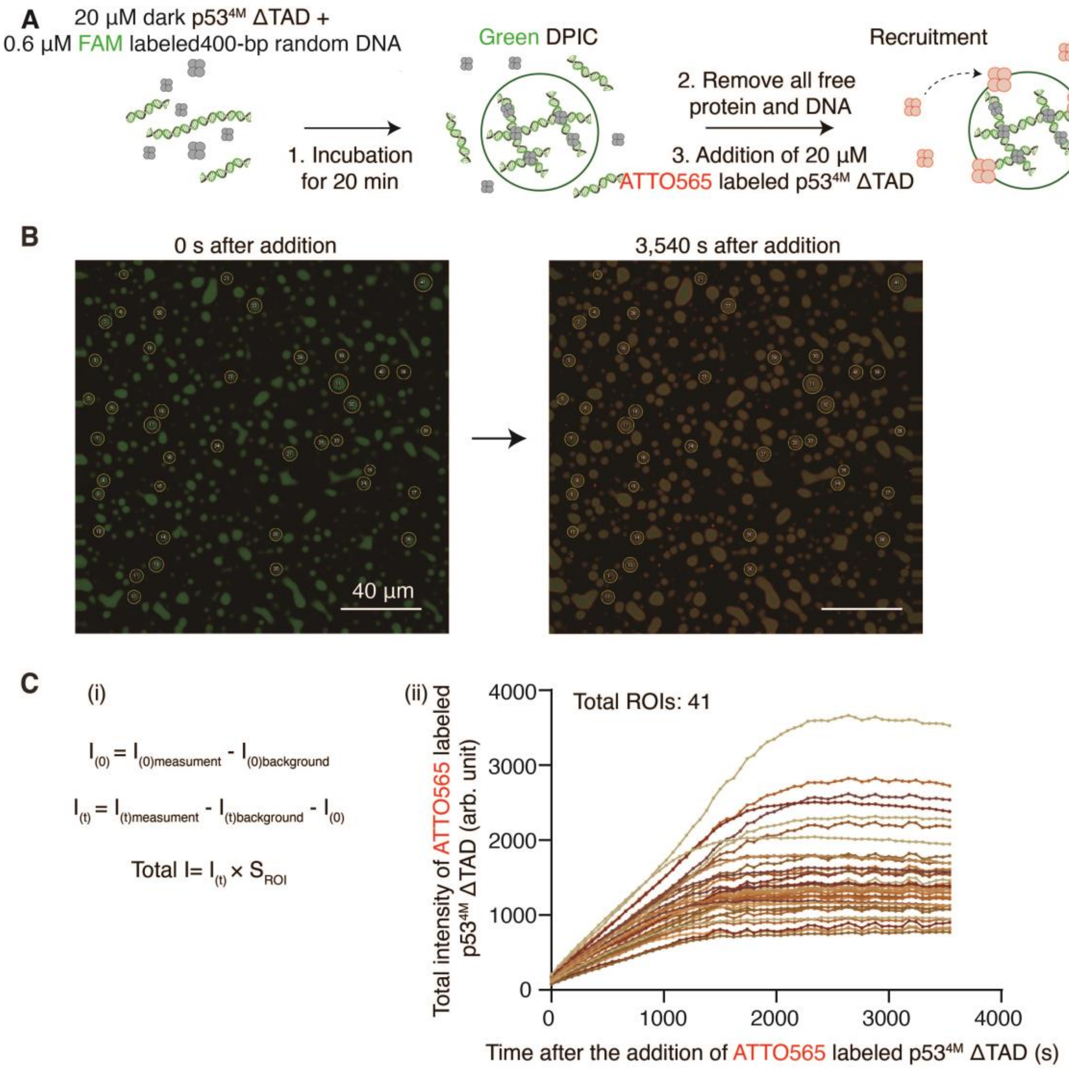


**Supplementary Fig. 3. The recruitment experiment in Fig. 1B.** (**A**) Schematic of the designed "recruitment experiment". (**B**) Imaging results of the recruitment experiment at 0 s and 3,540 s after the addition of 20 μM ATTO565-labeled p53$^{4M}$ ΔTAD. The green color represented the FAM-labeled 400-bp random dsDNA, and the red color represented the ATTO565-labeled p53$^{4M}$ ΔTAD. The yellow circles showed the measured DPICs. *In vitro* droplet experiments were conducted in a working buffer of 8 mM Tris-HCl (pH 7.5), 120 mM NaCl, 4% glycerol, and 16 mM DTT, without any crowding agents. The scale bar was 40 μm. (**C**) (i) The calculation of the recruitment of p53$^{4M}$ ΔTAD. $I_{(t)measurement}$ represented the average intensity of the p53$^{4M}$ ΔTAD in the yellow circles in (B) at each time point. $I_{(t)background}$ represented the average intensity of the surrounding environment at each time point. $I_{(0)}$ corresponded to the average protein intensity at 0 s after addition. And $S_{(ROI)}$ was the area of each yellow circle in (B). (ii) The growth curve of the total intensity of p53$^{4M}$ ΔTAD over time. Each curve represented one independent DPIC in the yellow circles in (B).

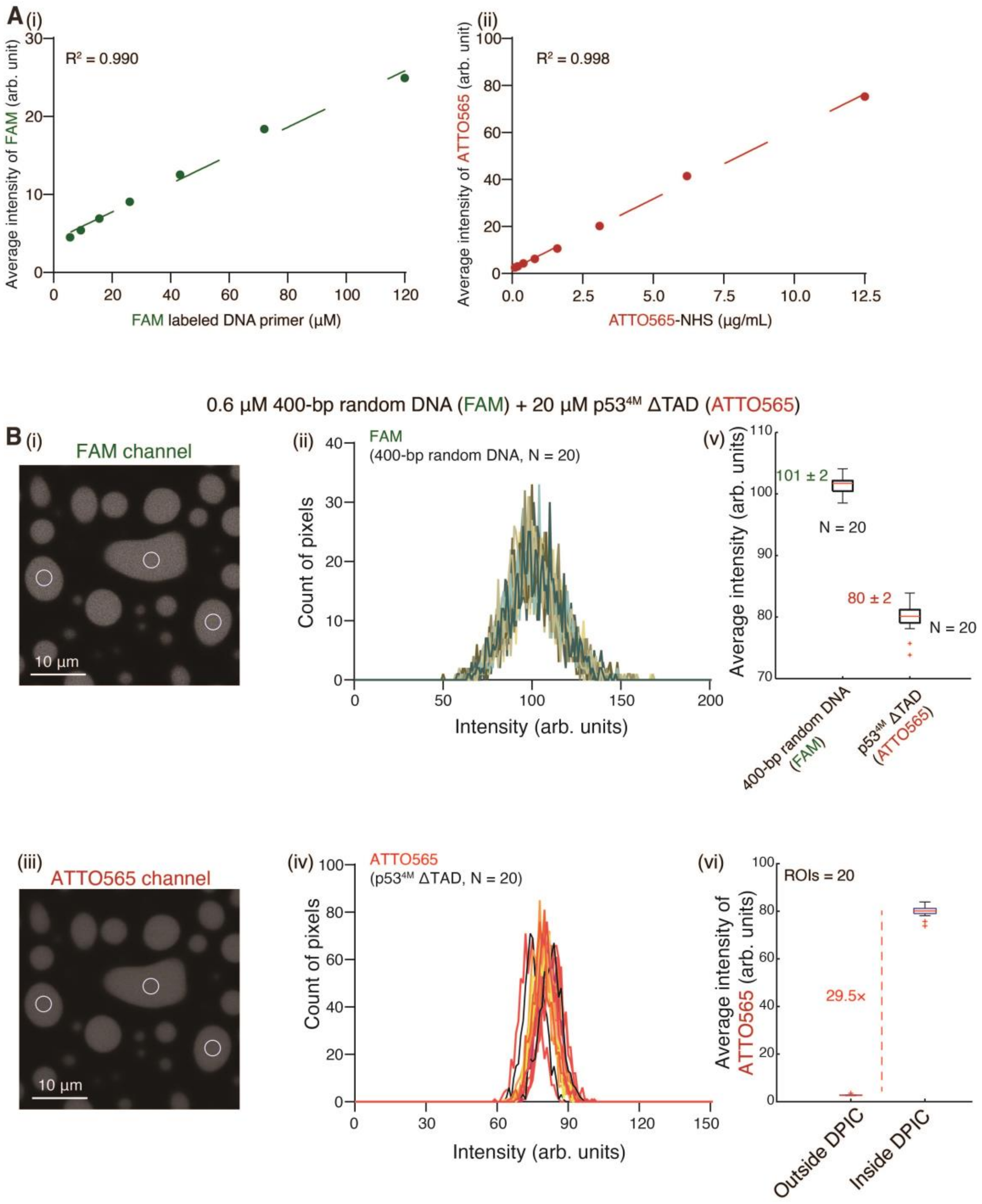


**Supplementary Fig. 4. Fluorescence distribution of p53$^{4M}$ ΔTAD and 400-bp random dsDNA inside and outside the DPIC.** (**A**) The relationship between the average intensity and the concentration of fluorescent molecules. (i) The FAM intensity and FAM-labeled dsDNA primer. (ii) The ATTO565 intensity and ATTO565-NHS. The dash lines represented the linear fit of the data, and the values of $R^2$ were shown. (**B**) Measurement of the fluorescent intensity of dsDNA and protein within DPIC. The experimental condition here is 0.6 μM FAM labeled 400-bp random dsDNA and 20 μM ATTO565 labeled p53$^{4M}$

ΔTAD. (i) The FAM channel. (ii) Distribution of FAM intensity of each pixel in the measurement regions. (iii) The ATTO565 channel. (iv) Distribution of ATTO565 intensity of each pixel in the measurement regions. The white circles represent three regions that were measured. Each curve corresponds to an independent measurement region. 20 regions were measured (N = 20). (v) The average intensity of FAM and ATTO565 of 20 independent measurement regions (N = 20). (vi) Comparison of the average intensity of ATTO565 inside and outside the DPIC. The intensity ratio was shown. For the boxplot, the red bar represents median. The bottom edge of the box represents $25^{th}$ percentiles, and the top is $75^{th}$ percentiles. Most extreme data points are covered by the whiskers except outliers. The '+' symbol is used to represent the outliers.

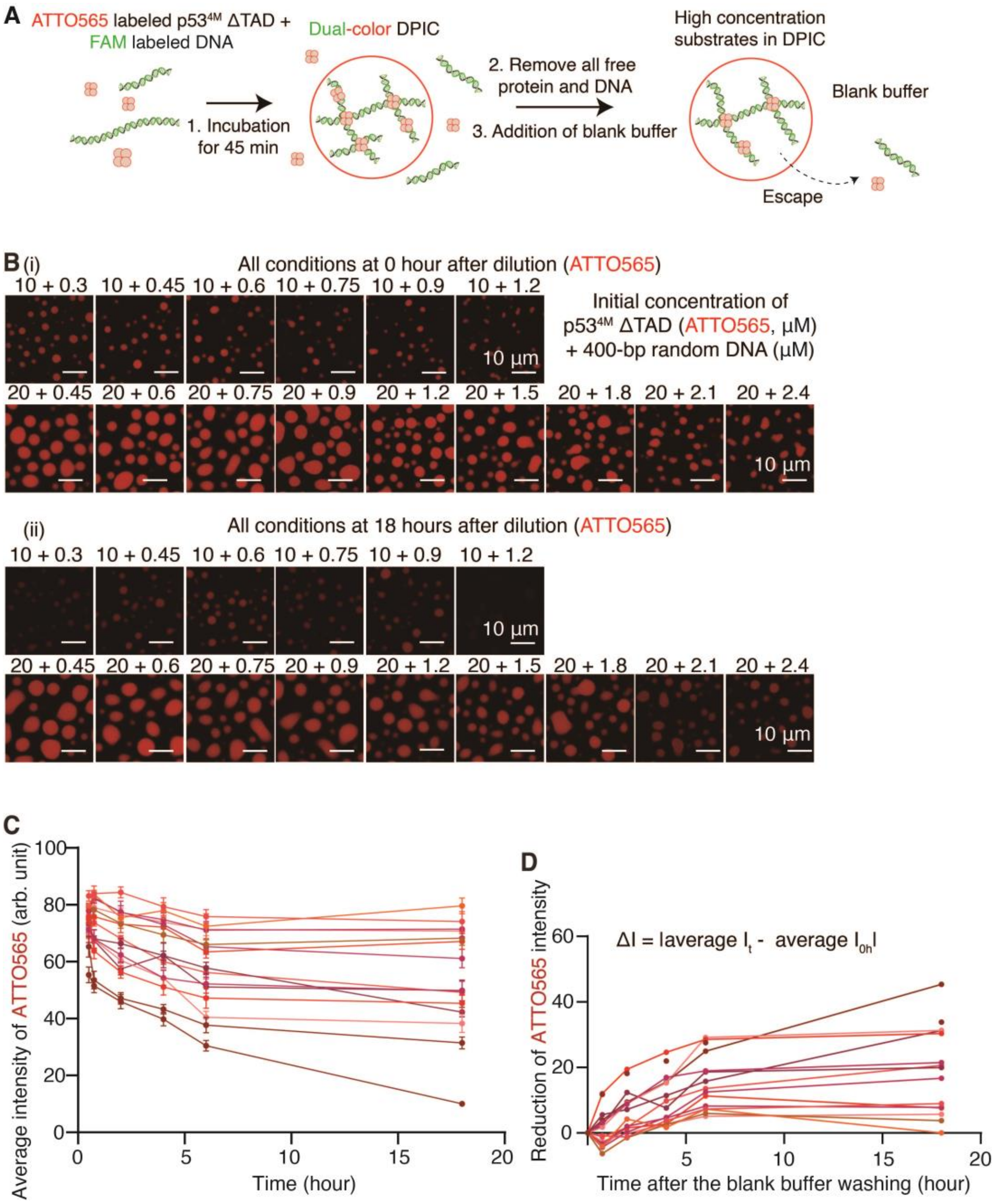


**Supplementary Fig. 5. The escape experiment in Fig. 1C.** (**A**) Schematic of the designed "escape experiment". (**B**) Imaging results of the escape experiments in 15 different experimental conditions at 0 hour (i) and 18 hours (ii) after the dilution by a blank buffer. The number above each image was the initial concentration of p53$^{4M}$ ΔTAD and 400-bp random dsDNA, respectively. The red color represented the ATTO565-labeled p53$^{4M}$ ΔTAD. *In vitro* droplet experiments were conducted in a working buffer of 8 mM Tris-HCl (pH 7.5), 120 mM NaCl, 4% glycerol, and 16 mM DTT, without any crowding agents. The scale bar was 10 μm. (**C**) The change of the average intensity of p53$^{4M}$ ΔTAD over time. Each curve represented one condition shown in (B). Each point represented

10 independent DPICs in one condition (N = 10). (**D**) The growth curve of the intensity reduction of p53$^{4M}$ ΔTAD over time. The intensity reduction was calculated by subtracting the average intensity of the protein inside the DPIC at each time point from the average intensity of the protein at 0 hour after dilution. Each curve represented one condition shown in (B).

#### 4. Supplementary Tables

**Supplementary Table 1. All simulation parameters were used in the mass-conserving dynamical model.**

| Symbol | Definition | Value | Units | Simulation value | Source |
|---|---|---|---|---|---|
| $\boldsymbol{u_0}$ | Concentration of dsDNA-bound protein at t = 0 s | $random$ $(0\sim0.5)$ | $\mu mol/L$ | 0.5 | |
| $\boldsymbol{v_0}$ | Concentration of free protein at t = 0 s | 30 | $\mu mol/L$ | varying | |
| $\boldsymbol{k_d}$ | Dissociation rate constant | 0.40 ± 0.13 | $s^{-1}$ | 0.36 | (*49*) |
| $\boldsymbol{\rho}$ | dsDNA length or sequence response to stability of arrest protein | [0.5, ∞] | dimensionless | 5.0 | Calibrated<br>In this study, set as the default value |
| $\boldsymbol{D_u}$ | Diffusion coefficient of dsDNA-bounded protein | $5.1\ \times\ 10^{-5}\boldsymbol{D_v}$ | $\mu m^2 s^{-1}$ | $0.05\boldsymbol{D_v}^*$ | Experiment |
| $\boldsymbol{D_v}$ | Diffusion coefficient of free protein | 18.0 ± 1.1 | $\mu m^2 s^{-1}$ | 0.17 | Our FCS data in Supplementary Fig. 2B(iii) |
| | | 15.4 ± 5.6 | $\mu m^2 s^{-1}$ | 0.17 | (*49*) |
| $\boldsymbol{k_1}$ | Recruitment rate coefficient | 1.35 ± 0.05 | $(\mu mol/L)^{-1}h^{-1}$ | 1.35 | Supplementary Table S2 |
| $\boldsymbol{\tilde{b}}$ | Self-triggered rate(intensity) | 93 ± 56 | $h^{-1}$ | 60 | Supplementary Table S2 |
| $\boldsymbol{k_3}$ | Conversion factor from fluorescence to concentration | 1.0~10.0 | dimensionless | 6.7 | Supplementary Fig. 4B(vi) |
| $\boldsymbol{b}$ | Self-triggered rate | $\frac{\tilde{b}}{k_3}$ | $h^{-1}$ | 8.95 | Fig. 1B |
| $\boldsymbol{k_2}$ | Maximal reduction of intensity per hour | 18.0 | $h^{-1}$ | 18 | Estimated<br>In Supplementary Fig. 5D |
| $\boldsymbol{\beta}$ | Half-saturation constant | 1.6 ± 0.4 | $\mu mol/L$ | 2.0 | Experiment |

* 0.05 is used for speeding up the numerical simulation.

**Supplementary Table 2. Model selections for recruitment response curve and escape response curve based on AIC and BIC criteria.**

| Model selection for $\tilde{f}(\tilde{u})$ | | |
|---|---|---|
| **Model (formula)** | **AIC** | **BIC** |
| $\tilde{f}(\tilde{u}) = a\tilde{u} + b$ | 520 | 524 |
| $\tilde{f}(\tilde{u}) = a\tilde{u}^2 + b\tilde{u} + c$ | 522 | 528 |
| $\tilde{f}(\tilde{u}) = a\ ln\tilde{u} + b$ | 551 | 555 |
| $\tilde{f}(\tilde{u}) = \frac{a\tilde{u}}{\tilde{u}+b}$ | 521 | 525 |

| Model selection for $\tilde{g}(\tilde{u},\tilde{v})$ | | |
|---|---|---|
| **Model (formula)** | **AIC** | **BIC** |
| $\tilde{g}(\tilde{u},\tilde{v}) = a(\tilde{u}+\tilde{v}) + b$ | -20 | -19 |
| $\tilde{g}(\tilde{u},\tilde{v}) = a(\tilde{u}+\tilde{v})^2 + b(\tilde{u}+\tilde{v}) + c$ | -23 | -21 |
| $\tilde{g}(\tilde{u},\tilde{v}) = aln(\tilde{u}+\tilde{v}) + b$ | -27 | -25 |
| $\tilde{g}(\tilde{u},\tilde{v}) = \frac{a}{(\tilde{u}+\tilde{v})+b}$ | -30 | -29 |

#### 1. Supplementary Movie Legends

**Supplementary Movie 1. 10 μM p53$^{4M}$ ΔTAD and 0.3 μM 400-bp random dsDNA formed the droplet-like DPICs.** The video was acquired for 2 hours after a 2-min incubation. 30-s shutter time was used. In this experiment, the dsDNA was labeled by FAM. The movie showed the merged signal of FAM with the bright field.

**Supplementary Movie 2. Numerical simulation of DPIC formation with $V_0$ = 20 and $\rho$ = 4.0.** The simulation time was set as 10000 seconds, with a time step of 0.02 seconds. Images were saved with a time array containing 200 points evenly distributed on a logarithmic scale, i.e. 200 images. Subsequently, we employed the IPython.display module along with the matplotlib.animation module in Python to create an animation, which was saved as an MP4 video file.

**Supplementary Movie 3. Numerical simulation of DPIC formation with $V_0$ = 25 and $\rho$ = 15.0.** All other parameters are the same to Supplementary Movie 2, except the $V_0$ and $\rho$ were set as 25 and 15.0.

**Supplementary Movie 4. Numerical simulation of DPIC formation with $V_0$ = 30 and $\rho$ = 22.5.** All other parameters are the same to Supplementary Movie 2, except the $V_0$ and $\rho$ were set as 30 and 22.5.

**Supplementary Movie 5. 20 μM p53$^{4M}$ ΔTAD and 0.6 μM 400-bp random dsDNA formed the droplet-like DPICs.** The video was acquired for 2 hours after a 2-min incubation. 30-s shutter time was used. In this experiment, the dsDNA was labeled by FAM. The movie showed the merged signal of FAM with the bright field.
